# Mutant forms of DDX3X with diminished catalysis form hollow condensates that exhibit sex-specific regulation

**DOI:** 10.1101/2023.03.19.533240

**Authors:** Michael C. Owens, Hui Shen, Amber Yanas, Maria Saraí Mendoza-Figueroa, Ellen Lavorando, Xiaoyu Wei, Him Shweta, Hsin-Yao Tang, Yale E. Goldman, Kathy Fange Liu

## Abstract

Mutations in the RNA helicase DDX3X, implicated in various cancers and neurodevelopmental disorders, often impair RNA unwinding and translation. However, the mechanisms underlying this impairment and the differential interactions of DDX3X mutants with wild-type (WT) X-linked DDX3X and Y-linked homolog DDX3Y remain elusive. This study reveals that specific DDX3X mutants more frequently found in disease form distinct hollow condensates in cells. Using a combined structural, biochemical, and single-molecule microscopy study, we show that reduced ATPase and RNA release activities contribute to condensate formation and the catalytic deficits result from inhibiting the catalytic cycle at multiple steps. Proteomic investigations further demonstrate that these hollow condensates sequester WT DDX3X/DDX3Y and other proteins crucial for diverse signaling pathways. WT DDX3X enhances the dynamics of heterogeneous mutant/WT hollow condensates more effectively than DDX3Y. These findings offer valuable insights into the catalytic defects of specific DDX3X mutants and their differential interactions with wild-type DDX3X and DDX3Y, potentially explaining sex biases in disease.

## Introduction

The DEAD-box RNA helicase DDX3X is a crucial component of numerous cellular and developmental processes^1^ that is conserved from yeast to humans^2^. Whole-body knockout of *Ddx3x* leads to embryonic lethality in mice^3^, and knockdown of Ddx3x in the developing murine nervous system disrupts cortex development^4^, highlighting the essential nature of DDX3X. DDX3X contains a helicase core common to all DEAD-box proteins that is flanked by two intrinsically disordered regions (IDRs) (**Fig. 1a**)^5^. The non-processive RNA unwinding activity of the helicase core helps DDX3X facilitate translation initiation for specific transcripts via unwinding their highly structured 5’ untranslated regions (5’UTRs) under unstressed conditions^6^. Upon cellular stress, the IDRs of DDX3X (particularly its N-terminal IDR) drive it to undergo liquid-liquid phase separation (LLPS) and participate in stress granule formation, which leads to translation repression of sequestered transcripts^7^.

**Fig. 1:**
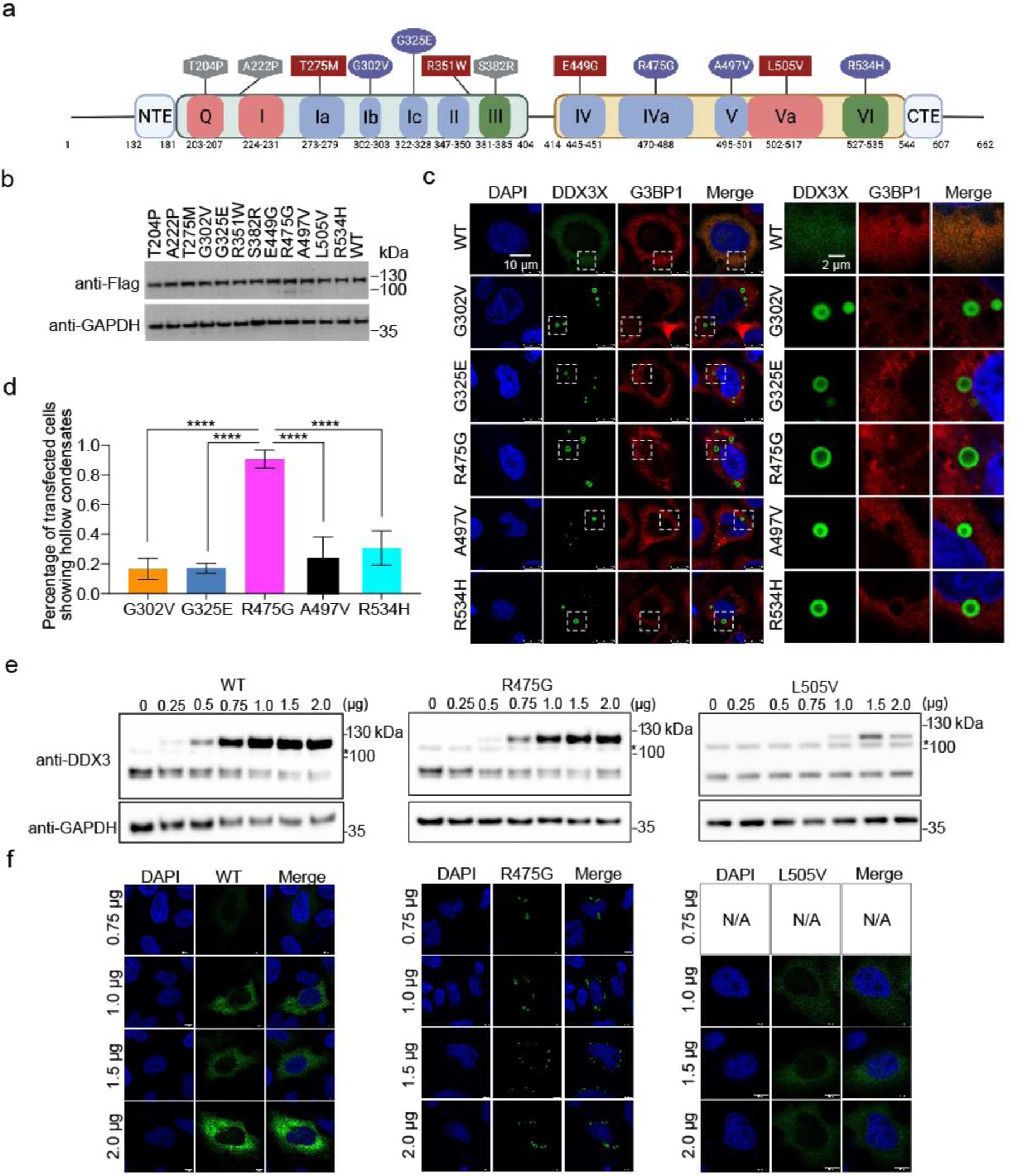
A subset of DDX3X disease mutants form unique hollow condensates in cells. **a**, Schematic of the conserved domains and motifs of DDX3X and the locations of each disease variant used in this study. Motifs responsible for ATP binding and hydrolysis (Q, I, and Va) are in red, RNA binding (Ia, Ib, Ic, II, IV, IVa, V) in blue, and coordination of ATPase and RNA binding (III and VI) in green. Diffuse variants are shown in squares, speckled variants in hexagons, and hollow mutants in ovals. N-terminal IDR = a.a. 1-131, N-terminal extension (NTE) = a.a. 132-181, N-terminal RecA-like domain = a.a. 182-403, linker = a.a. 405-413, C-terminal RecA-like domain = a.a.414-543, C-terminal extension (CTE) = a.a. 545-607, C-terminal IDR = 608-662. **b**, Western blots showing that the mClover3-tagged DDX3X variants were expressed at similar levels in HeLa cells. **c**, Representative images showing the localization of mClover3-tagged WT DDX3X or the indicated variants and mCherry-tagged G3BP1 in HeLa cells. The white boxes indicate regions zoomed on the right. Scale bars, 10 µm and 2 μm. **d**, Quantified percentage of transfected HeLa cells containing DDX3X variant hollow condensates. Values represent means ± s.d. from three independent measurements of 25 cells each. Significance was calculated using a two-tailed t-test. \*\*\*\**p* <0.0001. **e**, Western blot showing expression of exogenous mClover3-tagged WT DDX3X, L505V, and R475G and endogenous WT DDX3X in HeLa cells. * indicates a non-specific band produced by this antibody. **f**, Immunofluorescence imaging showing cellular distribution patterns of WT DDX3X, L505V, and R475G at the indicated plasmid concentrations **e**. Scale bar, 10 µm.

Recent studies have reported over 100 recurrent mutations in DDX3X across numerous cancers including medulloblastoma^8–10^, Burkitt’s lymphoma^11^, chronic lymphocytic leukemia and natural killer-T cell lymphoma^12–14^. Mutations in DDX3X are also linked to DDX3X syndrome, which accounts for 1-3% of otherwise unexplained intellectual disability cases^4,15–17^. These mutations often cause defects in the enzymatic activities (RNA binding, ATPase activity, and RNA unwinding), although the degree varies among pathogenic mutants^4,10,18^. As a consequence of disrupting the catalytic activity, several mutants of DDX3X are known to negatively impact translation^10^, in particular the translation of known DDX3X targets^4^. Despite the fact that the majority of these mutations are in the catalytic core of DDX3X and not in its IDRs, disease mutants of DDX3X frequently aberrantly phase separate in cells without addition of an external stressor^4,10^. DDX3Y, the sexually dimorphic homolog of DDX3X, is both a weaker ATPase and a stronger phase separator than DDX3X^19^. In concurrent work, we found that RNA plus WT DDX3X or DDX3Y form nano-sized RNA protein clusters (RPCs) with numerous copies of the proteins that optimally enhance helicase activity and may be nuclei for phase separation^20^. These results suggest that the enzymatic and phase separation properties of DDX3X and DDX3Y are intertwined, although much work remains to decipher how these defects contribute to the development of the corresponding disorders.

The problems associated with DDX3X mutants, as well as a wide range of other diseases, often manifest in a sex-biased manner. For example, DDX3X syndrome is significantly more common in XX individuals than in XY individuals^4^, while XX individuals with pediatric medulloblastoma have a higher survival probability than XY individuals^21^. Recently, a growing body of evidence suggests that some of these sex biases may stem from the different biochemical and biophysical properties of sexually dimorphic protein homologs^19,22–24^. DDX3X and its Y chromosome-encoded homolog, DDX3Y, are two of these sexually dimorphic homologs. Because mutants of DDX3X are often heterozygous^25,26^, the mutant allele will almost always be co-expressed with either wild-type (WT) DDX3X (in XX individuals) or with DDX3Y (in XY individuals). However, it is not known whether the enzymatic and condensation defects of DDX3X disease mutants are differently altered by co-expression with WT DDX3X or DDX3Y.

Here, we focused on understanding how a range of DDX3X disease mutants impact enzyme catalysis, condensation properties, and interactions with WT DDX3X or DDX3Y in cells. The results show that a specific subset of these mutants that occur more frequently across several cancers and DDX3X-related neurodevelopmental disorders than other mutants promote the formation of prominent condensate structures in cells. These mutants sequester WT DDX3X, DDX3Y, and additional proteins essential for various cellular functions, possibly explaining their more severe pathogenic outcomes. Our study also reveals the mechanism underlying formation of condensate structures under non-stressed conditions. These observations provide insights into the molecular basis of DDX3X-related pathogenesis in humans and, importantly, begin to reveal DDX3X/3Y-based differences in human health and disease.

## Results

### DDX3X disease mutants present with distinct cellular distribution patterns

Like other members of the DEAD-box family, the catalytic core of DDX3X consists of two RecA-like domains [amino acids (a.a.) 182 – 403 and 414 – 544] flanked by N- and C-terminal intrinsically disordered regions (IDRs) (a.a. 1 – 131 and 608 – 662). Within the two RecA-like domains are twelve conserved motifs that have been implicated in either binding and hydrolyzing ATP (Q, I, and Va motifs, in red), binding RNA (Ia, Ib, Ic, II, IV, IVa, and V motifs, in blue) or coordinating these functions (III and VI motifs, in green)^5^ (**Fig. 1a**). In this work, we studied one disease mutation in (or near) each of these twelve conserved motifs in an effort to investigate the effects of mutants across the DDX3 catalytic core.

To begin, we expressed either WT DDX3X or one of twelve mutants (each tagged with N-terminal Flag and C-terminal mClover3) (**Fig. 1a**) to similar levels in Henrietta Lacks’ (HeLa) cells (**Fig. 1b**). WT DDX3X showed diffuse localization throughout the cytoplasm in unstressed cells, which is consistent with previous studies^4,10,19^ (**Fig. 1c**). Strikingly, five of the mutants (G302V, G325E, R475G, A497V, and R534H) formed spherical cytoplasmic puncta in which DDX3X was enriched in the outer shell but depleted in the center (**Fig. 1c**). Hereafter, we refer to these mutants as the “hollow” mutants. The percentages of transfected cells with any hollow puncta varied among these five hollow mutants; while four of these mutants formed hollow puncta in ∼20% of cells, R475G formed hollow puncta in greater than 90% of cells (**Fig. 1d**). R475G has previously been linked to severe neurodevelopmental phenotypes^4^, suggesting disease severity may be correlated with this high propensity for forming hollow puncta. This morphology was also observed in HEK293T cells for all hollow mutants (**Extended Data Fig. 1a**). Orthogonal projections of R475G further support the hollow structure of the puncta with the mutant protein enriched in the shells (**Extended Data Fig. 1b**). To ensure that hollow morphology was not due to overexpression, we titrated the R475G-mClover3 expression plasmid and quantified protein expression via Western blot. R475G hollow puncta were observed at the same concentrations as endogenous WT with an increasing number of hollow condensates per cell with increasing plasmid concentration (**Fig. 1e and 1f**). Across this concentration range, both WT and the diffuse mutant L505V remained diffuse throughout the cytoplasm (**Fig. 1e and 1f**). Additionally, expressing Flag-tagged R475G (lacking mClover3) also formed hollow puncta in cells, indicating that the fluorescent tag did not influence the morphology (**Extended Data Fig. 1c**).

Four of the other mutants used in this study (T275M, R351W, E449G, and L505V) were diffuse in the cytoplasm like WT DDX3X; these mutants are hereafter referred to as “diffuse” mutants (**Extended Data Fig. 1d**). Additionally, we observed that the remaining three of the mutants (T204P, A222P, and S382R) presented with small foci with irregular shape throughout the cytoplasm; these mutants are hereafter referred to as “speckled” mutants (**Extended Data Fig. 1d**).

Upon cellular stress, WT DDX3X undergoes LLPS and enters stress granules (SGs)^7^. This led us to consider whether the hollow puncta we observed were, in fact, SGs. Under unstressed conditions, none of the mutants’ puncta colocalized with the canonical SG marker G3BP1^27^ (**Fig. 1d and Extended Data Fig. 1d**). Upon arsenite treatment, all but the speckled DDX3X mutants colocalized with G3BP1, suggesting that they entered SGs (**Extended Data Fig. 1e**). Collectively, these results suggest that DDX3X mutants have distinct cellular condensation propensities, expression patterns, and possibly different protein interactomes compared to WT DDX3X.

### DDX3X hollow mutants display decreased ATPase and RNA release activities

To test how the DDX3X mutants disrupt enzyme catalysis, we purified WT DDX3X and each mutant (**Extended Data Fig. 2a**). While we were able to purify all the diffuse and hollow mutants, the speckle mutants always aggregated during purification attempts. The three speckled mutations (T204P, A222P, and S382R) all sit in the N-terminal RecA-like domain. In the apo state structure (PDB 5E7I) (**Fig. 2a, top**), Thr204 is at the N-terminus of an alpha helix, flanked by two prolines. T204P thus results in a stretch of three proline residues in a row. Ala222 and Ser382 are both in the central beta sheet that runs through the middle of the N-terminal RecA-like domain, and the backbone carbonyl of Ala222 hydrogen bonds with the backbone amine of Ser382. The A222P and S382R mutations individually may disrupt the formation of this beta sheet, causing instability in the protein. When expressed in HeLa cells, the speckled mutants precipitated to the cell pellet after lysis using radioimmunoprecipitation assay (RIPA) buffer (**Fig. 2a, bottom**). This apparent instability, together with the resistance to purification of these three mutants implies that these mutations likely result in misfolding of the N-terminal RecA-like domain.

**Fig. 2:**
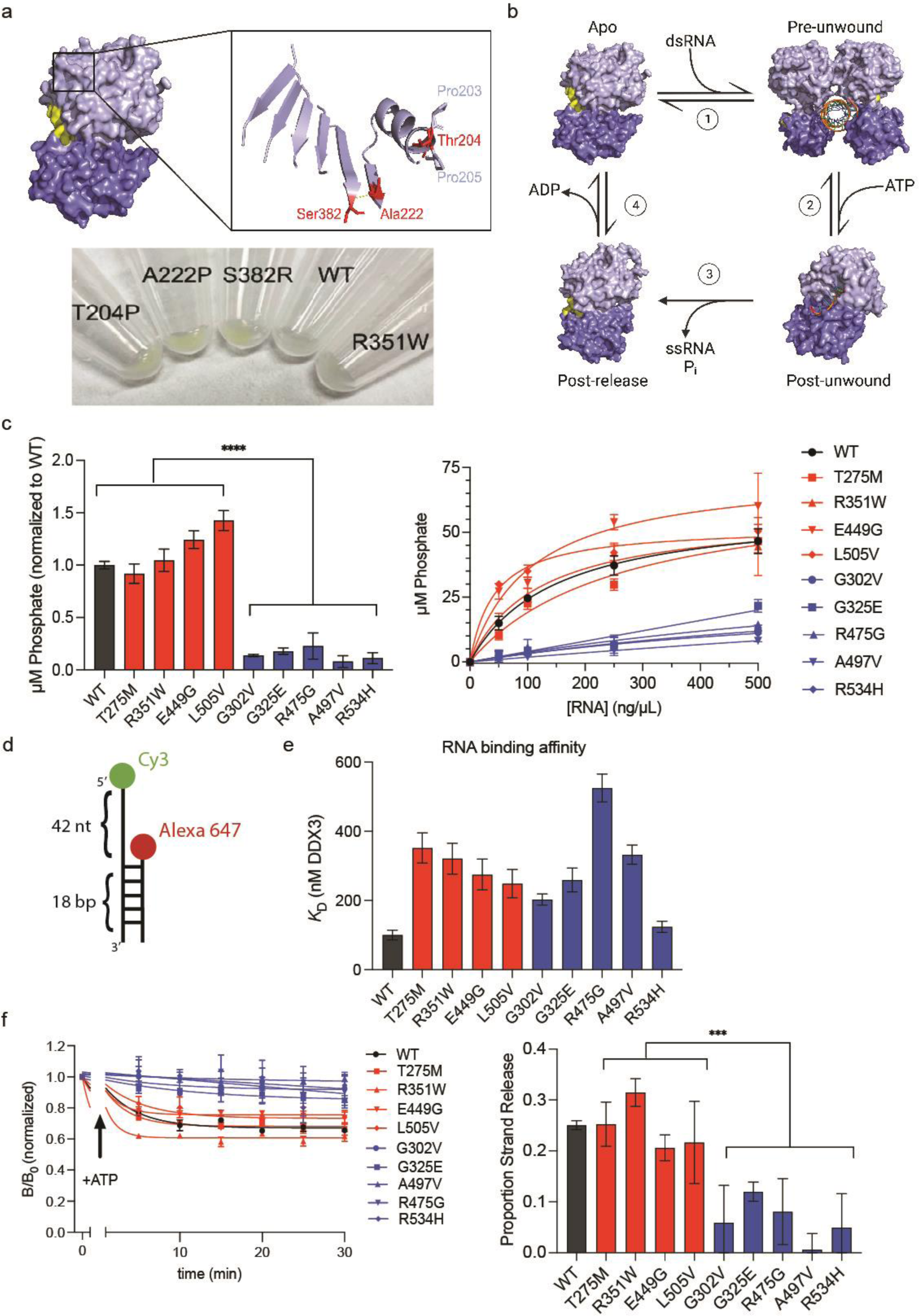
Hollow mutants are defined by their decreased ATPase and RNA strand release activities. **a**, Top, DDX3X apo structure zoomed into the three speckle mutants (in red). Pro203 and Pro205 flanking Thr204 are indicated in light blue. The yellow dashed line indicates the backbone hydrogen bond between Ala222 and Ser382. Bottom, pellets of HeLa cells expressing mClover3-tagged WT DDX3X or the indicated mutants, lysed in RIPA buffer. The yellow color of the three speckle mutant pellets indicates the relative insolubility of these mutants compared to WT DDX3X and R351W (a diffuse variant). **b,** Current model of the DDX3X-catalyzed helicase reaction. The N-terminal RecA-like domain is in light blue, the C-terminal RecA-like domain is in dark blue, and the RNA substrate is in orange. In the apo state (PDB 5E7I), DDX3X can bind dsRNA as a dimer (1) to form the pre-unwound state (PDB 6O5F). Upon addition of ATP, each monomer can perform an RNA strand separation (2), converting to the post-unwound state (PDB 2DB3). Following ATP hydrolysis, DDX3X releases both the inorganic phosphate and the product ssRNA strand (3), forming the post-release state (PDB 4PXA). Finally, ADP release (4) allows DDX3X to return to the apo state. **c,** Left, ATPase activity of mCherry-tagged WT DDX3X or the indicated mutants, measured after a 30-minute reaction in the presence of 100 ng/μL HEK293T total RNA and normalized to WT. Significance was calculated using a two-way ANOVA, \*\*\*\**p* < 0.0001. Values represent mean ± s.d., n = 3. Right, ATPase activity of mCherry-tagged WT DDX3X or the indicated measured at the indicated concentration of total RNA. Values were fit to Michaelis-Menten kinetics. **d,** Schematic of the dsRNA probe used for EMSA and fluorescence anisotropy experiments. **e,** Summarized *K_D_* values for the indicated construct, calculated via EMSA using mCherry-tagged protein in D. Values represent mean ± s.e.m., n = 3. **f**, Left, time courses tracing the B/B_0_ (or fraction anisotropic) short strand RNA over time in the presence of 1mM ATP and MBP-tagged WT DDX3X or the indicated mutants. Right, change in B/B_0_ signal (proportion strand release) from 0-30 minutes graphed for each protein.

The current model of the DDX3X catalytic cycle is as follows (**Fig. 2b**): apo DDX3X (DDX3X alone, PDB 5E7I)^28^ binds to dsRNA as a multimer (without IDRs, the crystallization construct forms a dimer, but evidence for higher order assemblies exist for full-length protein^20,29^) to adopt the pre-unwound state (PDB 6O5F)^30^. ATP binding produces the post-unwound state (represented in **Fig. 2b** by the related DEAD-box protein Vasa with ssRNA and AMPPNP, PDB 2DB3) ^31^. Upon ATP hydrolysis, DDX3X releases its single stranded product, adopting the post-release state (DDX3X with ADP, PDB 4PXA)^18^. ADP release completes the cycle (**Fig. 2b** and **Movie 1**). Given the crucial role of ATP hydrolysis and that the ATPase activity of DEAD-box enzymes often influences their condensation properties^32^, we assayed the ATP hydrolysis activity (**Fig. 2b**, **step 3**) of all the mutants we were able to purify using malachite green to detect phosphate release as previously described^19^ (**Fig. 2c, left**). When tested at equal protein concentration, all of the hollow mutants exhibited significantly lower ATPase activities than the diffuse mutants and WT DDX3X, suggesting that ATPase activity may correlate with their propensity to form hollow condensates in cells. Because the ATPase activity of DEAD-box proteins is RNA-stimulated, we questioned whether these differences in ATPase activity could be due to decreased RNA binding affinity (**Fig. 2b**, **step 1**). A mutant whose sole problem is a deficiency in RNA binding relative to WT is expected to reach the same maximum phosphate release rate as the WT at high RNA concentration. To test this, we performed electrophoretic mobility shift assays (EMSAs) with each mutant using the dsRNA illustrated in **Fig. 2d**. We found that all of the mutants, except for R534H, had RNA binding affinities weaker than WT, and that unlike ATPase activity, RNA binding did not correlate with the cellular morphology (**Fig. 2e** and **Extended Data Fig. 2b**). We then repeated the malachite green assay across several total RNA concentrations (**Fig. 2c, right**), finding that WT and the diffuse mutants were more active than the hollow mutants at all RNA concentrations. Additionally, the ATPase activity of WT and the diffuse mutants increased robustly with increasing RNA concentrations, while the hollow mutants displayed a much more modest increase.

Given that ATP hydrolysis triggers RNA strand release (**Fig. 2b**, **step 3**), we next studied whether the ATPase deficiency impairs strand release using a single-molecule fluorescence time-resolved anisotropy approach. Briefly, a dsRNA molecule labeled with Cy3 (long, overhang strand) on one strand and Alexa647 (short strand) on the other (**Fig. 2d**) was incubated with either WT DDX3X or the indicated mutant (**Extended Data Fig. 2c**). Since DDX3X is a non-progressive helicase, it must unbind and rebind RNA as it progresses through the enzymatic cycle. This activity can be monitored with fluorescence anisotropy as the short strand RNA is released (and tumbles more quickly) with the addition of ATP. This activity is interpreted as strand release activity from either the DDX3X enzyme and/or escape from RNA-protein clusters (catalytically active nano-sized RNA-protein assemblies) of DDX3X (see our concurrent manuscript^20^). Specifically, the fraction of anisotropic (slow tumbling) RNA (B/B_0_) was monitored for the different species of RNA (long single-stranded, short single-stranded, and duplex RNA) (**Fig. 2f, left**). The fraction of anisotropic RNA decreased for the short strand RNA over time for the WT DDX3X and diffuse mutants, however the fraction of anisotropic RNA for the short strand decreased significantly less for the hollow mutants (**Fig. 2f, right**). On the other hand, the fraction anisotropic RNA for the long strand and duplex species remained high with and without the addition of RNA for WT, diffuse, and hollow DDX3X, indicating stably bound protein (**Extended Data Fig. 2d** and 2e). Collectively, these results suggest that hollow mutants of DDX3X have severe defects in turnover involving ATP hydrolysis and RNA strand release, while diffuse mutants are comparable to WT in these activities.

### The decreased enzymatic activity of DDX3X hollow mutants is due to defects throughout the catalytic cycle

The diminished catalysis we observed in **Fig. 2** could be the result of difficulties progressing through any (or multiple steps) of the DDX3 catalytic cycle. To obtain a thorough mechanistic understanding of how individual mutations disrupt catalysis, we traced the interactions of each mutated residue throughout the DDX3X catalytic cycle. For example, Arg475 forms a salt bridge with Asp455 in the pre-unwound state which seems to position several residues for RNA binding (**Fig. 3a**). Flipping the residues of this salt bridge (D455R/R475D, **Extended Data Fig. 3a**) resulted in partial rescue of both ATPase activity (**Fig. 3b**) and RNA binding activity (**Extended Data Fig. 3b** and 3c), suggesting that the disruption in ATPase activity for R475G stems, at least in part, from disruption of RNA binding (**Fig. 2b, step 1**). However, restoration of the positive charge (R475K, **Extended Data Fig. 3a**) showed a more robust rescue of ATPase activity than the salt bridge swap (**Fig. 3b**), suggesting a role for a positive charge at position 475 later in the catalytic cycle and implying that R475G may disrupt several catalytic steps.

**Fig. 3:**
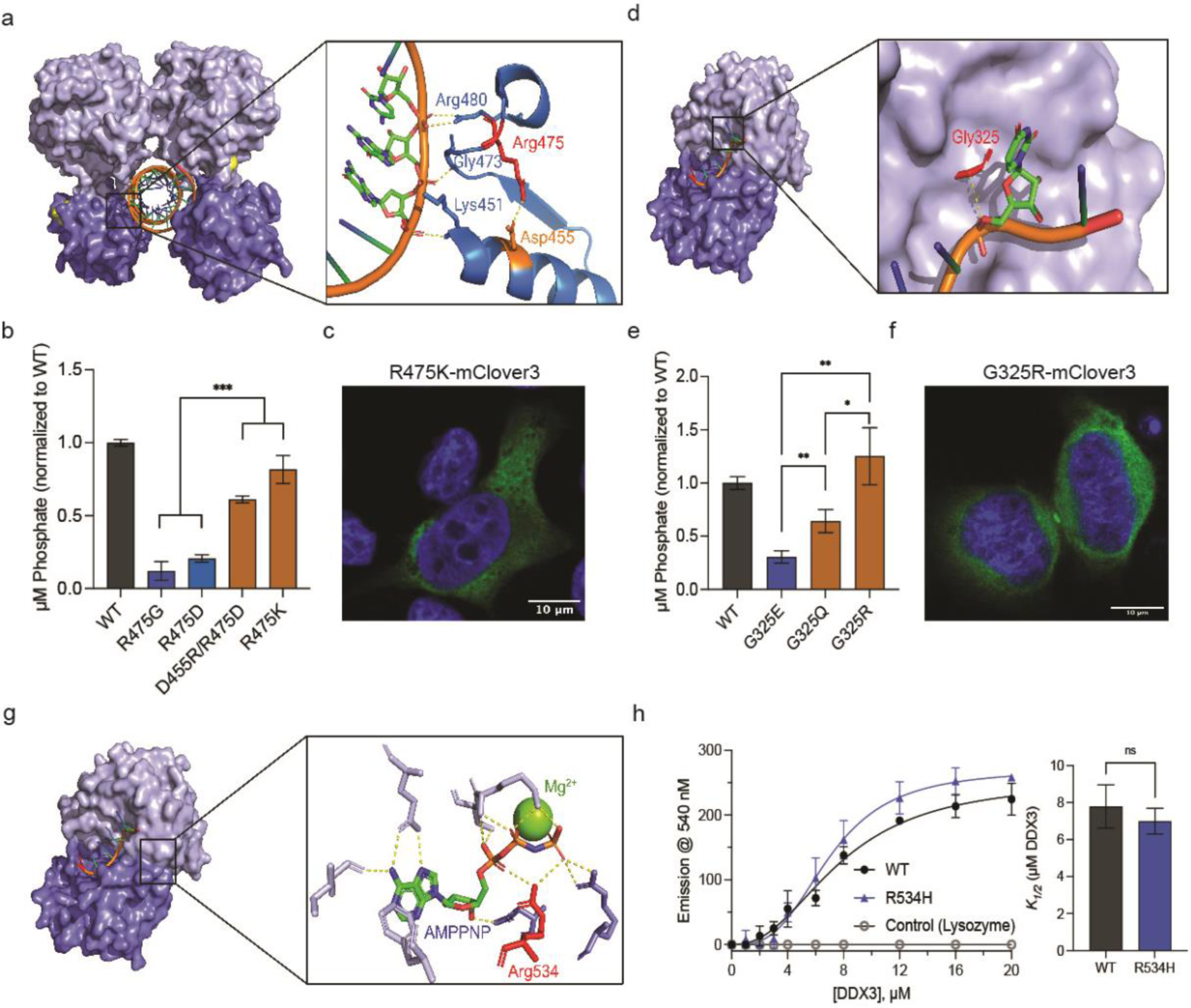
Mechanistic studies reveal how DDX3X mutants interfere with catalysis. **a,** Pre-unwound structure of DDX3X zoomed on Arg475 (red) and the surrounding RNA binding residues. Asp455 noted in orange, a putative salt bridge between these residues is indicated by the dashed yellow line. Interactions between Lys451, Gly473, and Arg480 and the RNA backbone are indicated by their respective yellow dashed lines. **b,** ATPase activity of mCherry-tagged WT DDX3X, R475G, R475D, D455R/R475D, and R475K normalized to WT. Significance was calculated using a two-tailed t-test. ***, *p*<0.001. Values represent mean ± s.d., n = 3. **c,** A representative image of R475K-mClover3 in HeLa cells. **d,** Post-unwound structure of DDX3X zoomed in on Gly325 (red, numbered according to DDX3X homology). The dashed line indicates the 3.6 Å distance between the alpha carbon of Gly325 and the RNA backbone. **e,** ATPase activity of mCherry-tagged WT DDX3X, G325E, G325Q, and G325R normalized to WT. Significance was calculated using a two-tailed t-test. **, *p*<0.01, *, *p*< 0.05. Values represent mean ± s.d., n = 3. **f,** A representative image of G325R-mClover3 in HeLa cells. **g,** Post-unwound structure of DDX3X zoomed in on ATP-binding pocket, showing residues involved in ATP binding and Mg^2+^ cation (yellow sphere). Arg534 (numbered with DDX3X homology) is indicated in red. Yellow dashed lines indicated interactions between residues and ATP. **h,** TNP-ATP binding curve for MBP-tagged WT DDX3X, R534H, and negative control (lysozyme). Right, bar graph showing *K_1/2_* values for WT and R534H. Significance was calculated using a two-tailed t test. ns, not significant. Values represent mean ± s.e.m., n = 3.

Gly325 sits in a positive cleft on the N-terminal RecA-like domain. In the pre-unwound state, this cleft is solvent accessible, but during the transition to the post-unwound state (**Fig. 2b, step 2**) this cleft accepts the newly unwound RNA (**Fig. 3d**), which we hypothesize is blocked by the G325E mutation. To evaluate whether G325E hinders catalysis due to steric or electrostatic effects, we generated DDX3X mutants G325Q and G325R and conducted further ATPase assays (**Extended Data Fig. 3a**). While G325Q (uncharged but as large as glutamate) partially rescued ATPase activity, G325R (positively charged and larger than glutamate) showed a full rescue (**Fig. 3e**). These results suggest that the negative charge introduced by the disease mutant G325E is a major disruptor of DDX3X catalysis. This is supported by time-resolved fluorescence anisotropy studies, as RNA mixed with G325R showed significantly higher change in the fraction of anisotropic molecules than RNA mixed with G325E, indicating a higher degree of strand release (and escape from RPC clusters), which is indicative of increased unwinding activity (**Fig. S3d**).

The Arg534 residue is located in the C-terminal RecA-like domain. In the post-unwound state, Arg534 coordinates the ATP’s α and Ɣ phosphates where it probably participates in hydrolysis (**Fig. 3g**). The R534H mutation most likely disrupts only the ATP hydrolysis step (**Fig. 2b, step 3**), as Arg534 is distal from the ATP binding site during all of the other states. Supporting this, R534H can bind both dsRNA (**Fig. 2e**) and the ATP analog TNP-ATP^33^ (**Fig. 3h**) as efficiently as WT DDX3X, indicating that R534H can form both the pre-unwound (dsRNA-bound) and post-unwound (ssRNA and ATP-bound) complexes as readily as WT DDX3X.

We next expressed R475K and G325R (tagged with C-terminal mClover3) in cells to assess whether the rescued ATP hydrolysis was concurrent with a loss of hollow morphology in cells (**Fig. 3c** and **3f**). Indeed, both rescue mutations restored the diffuse cytoplasmic localization present in WT DDX3X (hollow puncta were not observed). Given that ATP hydrolysis is necessary for the release of bound RNAs^18,28,31,34,35^, it is possible that the RNA trapped in the protein functions as a multivalent platform that recruits additional DDX3X molecules and other proteins, contributing to hollow puncta formation.

### RNA binding affinity influences inter- and intramolecular dynamics of DDX3X mutant hollow condensates

Unlike WT DDX3X, which only undergoes LLPS under stressed conditions^7^, select DDX3X mutants have increased phase separation propensities leading to puncta formation under unstressed conditions ^4,10^. Hollow puncta are likely formed via LLPS, as we were able to observe fusion of two hollow condensates while performing live-cell imaging (**Extended Data Fig. 4a**). In addition to fusion, a key trait of liquid-liquid phase separated condensates is that they are in a dynamic equilibrium with their surroundings^36^. To directly study the molecular exchange between the cytoplasm and the hollow condensates formed by DDX3X mutants, we performed “full bleach” fluorescence recovery after photobleaching (FRAP) experiments, in which we bleached an entire hollow condensate and measured the dynamics of fluorescence signal recovery (**Extended Data Fig. 4b, left**). We found that the fluorescence of G302V, G325E, A497V, and R534H hollow condensates recovered gradually, while the fluorescence of R475G hollow condensates recovered very little in the 100 s experimental timeframe (**Fig. 4a, left and 4b**, **top**). Additionally, “partial bleach” experiments were carried out to reveal the mobility of molecules within the hollow condensates (**Extended Data Fig. 4b, right**)^37^. Similarly, the fluorescence of R475G hollow condensates showed almost no recovery after partial bleaching unlike the other mutants, consistent with the full bleach results (**Fig. 4a**, **right and 4b**, **bottom**). The lower inter- and intramolecular dynamics of the R475G hollow condensates sets them apart from the other mutant hollow condensates. The uniquely low dynamics of R475G are in line with our previous finding that the percentage of cells expressing hollow condensates was much higher for R475G than for the other mutants (**Fig. 1d**). Previous studies suggest that RNA binding deficiencies of RNA-binding proteins such as FUS (fused in sarcoma) decreased the dynamics of the protein-RNA condensates^38^. While all the hollow mutants (save for R534H) displayed weaker RNA binding affinity than WT DDX3X, R475G had the weakest RNA affinity (**Fig. 2e**). This suggests that the decreased RNA binding of R475G may decrease the dynamics of its hollow condensates relative to the other hollow mutants.

**Fig. 4:**
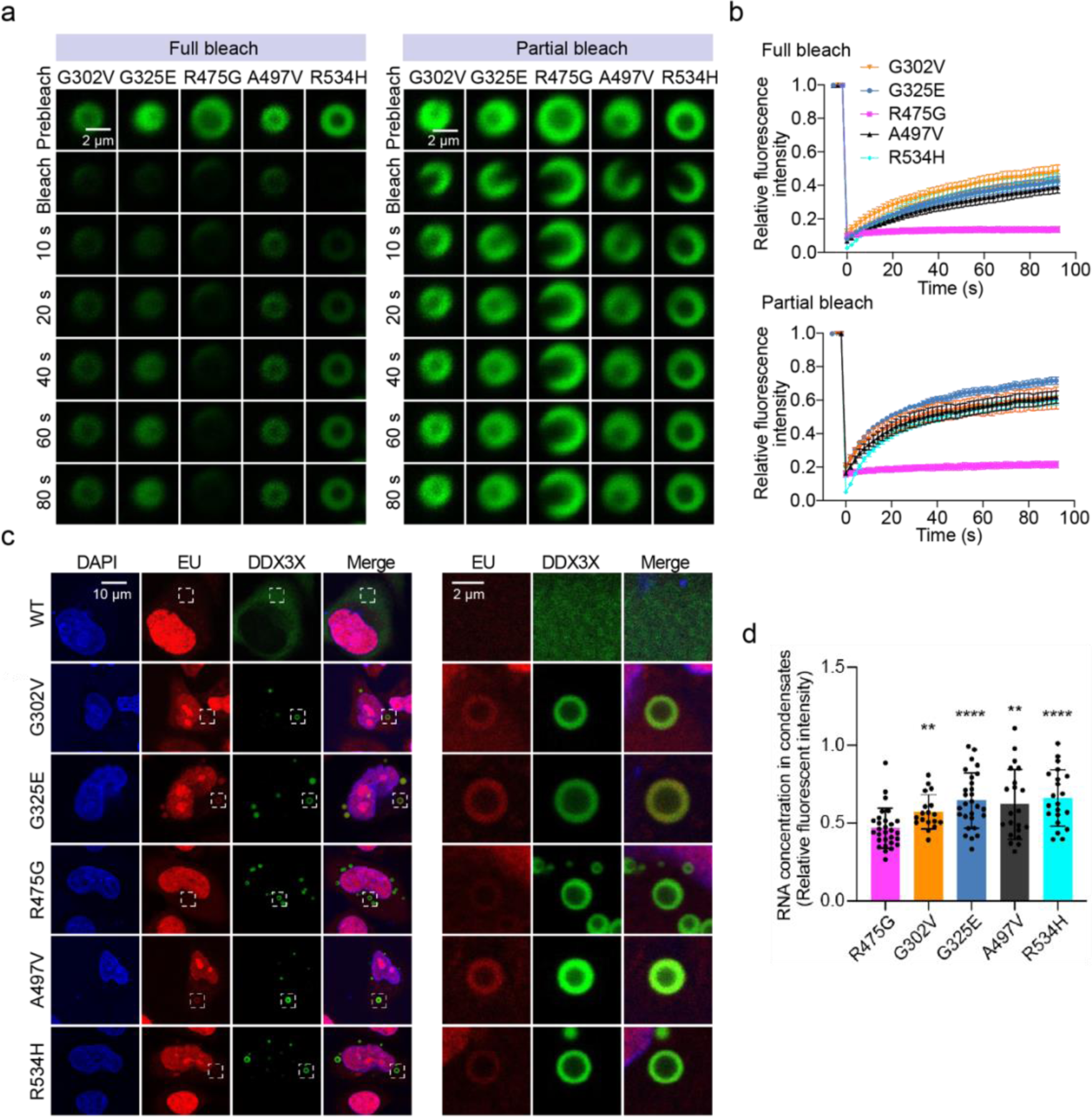
RNA-binding affinity differentiates the inter- and intramolecular dynamics of DDX3X hollow puncta. **a,** Time-lapsed images of representative FRAP experiments on hollow condensates of the indicated mClover3-tagged DDX3X variants expressed in HeLa cells. Full bleach (left) and partial bleach (right) experiments were performed to measure the inter- and intramolecular dynamics (respectively) of each condensate. Scale bar, 2 μm. **b,** Recovery curves for the indicated mClover3-tagged DDX3X variants in HeLa cells. Top: recovery plot for full bleach experiments. Bottom: recovery plot for partial bleach experiments. The traces of the FRAP data represent means ± s.e.m. from at least three biologically independent experiments. **c,** Left: imaging of EU-labeled RNAs (red) in HeLa cells expressing mClover3-tagged WT or DDX3X hollow variants. Scale bar, 10 μm. Right: zoomed-in view of the boxed area from the left figures. Scale bar, 2 μm. **d,** Quantification of EU fluorescence intensity for each of the hollow condensates represented in d. Values were normalized using the fluorescence intensity in the nucleoplasm of each cell. Values represent means ± s.d. from 20 cells. Significance was calculated between R475G and the other variants using a two-tailed t-test. \*\**p* <0.01; \*\*\*\**p* <0.0001.

Furthermore, to assess whether the decreased RNA binding affinities measured *in vitro* correspond to decreased RNA binding in cells, we utilized click chemistry to fluorescently label nascent RNA in HeLa cells after incorporation of 5-ethynyluridine (5-EU) (**Extended Data Fig. 4c**). The results showed that the cytoplasmic RNAs were enriched in the outer shells of hollow condensates relative to the cytoplasm and the center of the condensates (**Fig. 4c**). After quantifying the RNA signal intensity in the condensates’ shells of each of the hollow mutants (normalizing to the RNA signal in the nucleoplasm), we found that R475G hollow condensates had the weakest 5-EU signals of all the tested mutants (**Fig. 4d**), consistent with the *in vitro* EMSA data (**Fig. 2e**).

### R475G hollow condensates are enriched with proteins in various signaling pathways and lack translation machinery

Under unstressed conditions, DDX3X mutant hollow condensates lack the SG marker protein G3BP1, indicating that these condensates are not SGs (**Fig. 1c**). This raised the possibility that these condensates may sequester unique proteins and have functions divergent from those of SGs. To understand the potential biological functions of these hollow condensates, we sought to identify their protein components using an adapted ascorbate peroxidase (APEX2)-based proximity labeling method^39–41^ (**Fig. 5a**). The labeling efficiency and specificity of APEX2-tagged DDX3X were examined as previously shown^19^. In this study, we prioritized comparing the protein networks of R475G and WT DDX3X, as R475G displayed the highest propensity to form hollow condensates among all the mutants tested.

**Fig. 5:**
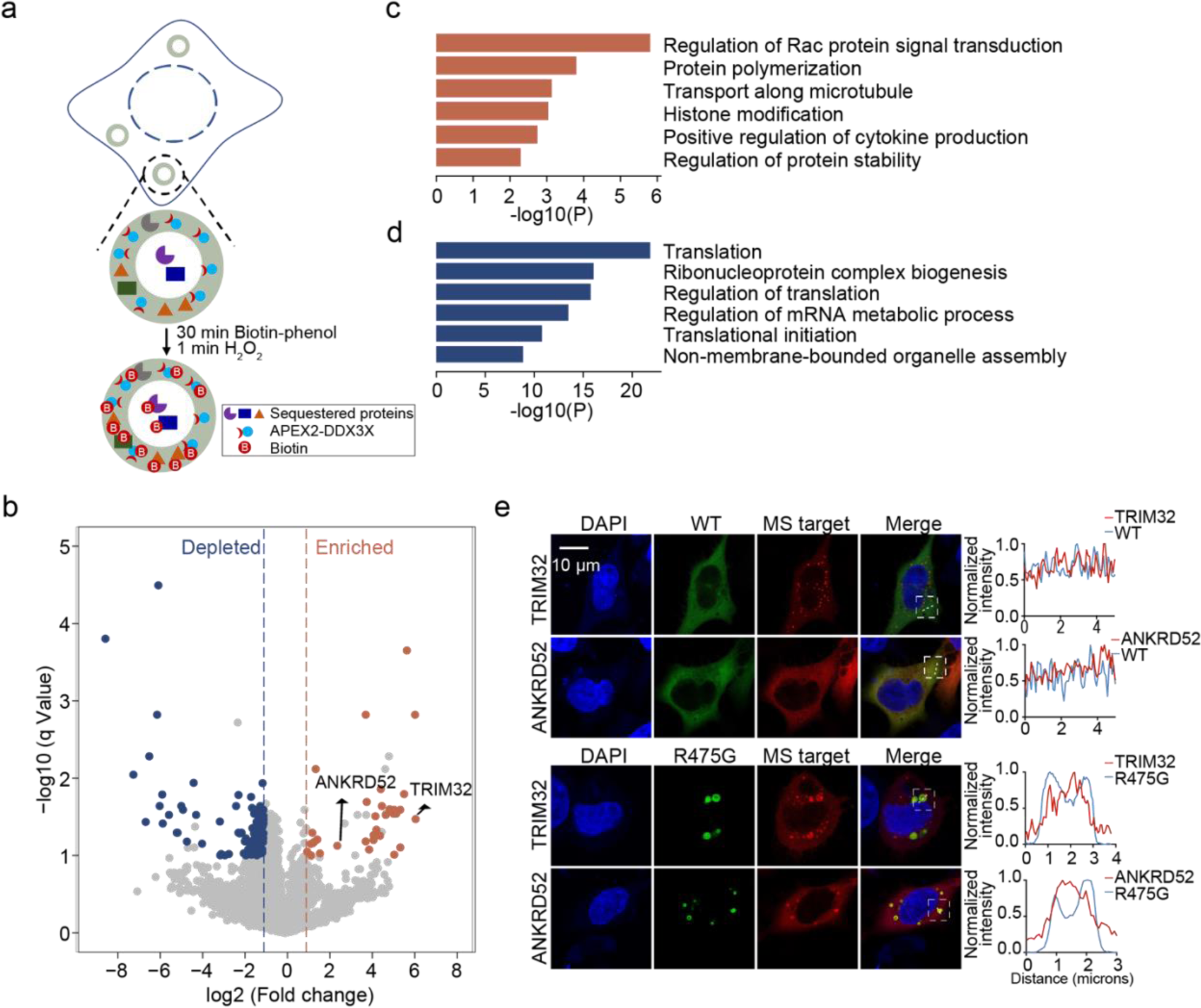
R475G hollow condensates enrich proteins in various signaling pathways and deplete translation machinery. **a,** Schematic of APEX2-mediated proximity labeling of proteins in R475G hollow condensates. **b,** Volcano plots showing differential protein enrichment via streptavidin pull-downs from APEX2-R475G compared to APEX2-WT expressing cells. Differentially enriched proteins are shown in orange (adjusted *p*-value < 0.1, log_2_ fold change > 1), and differentially depleted proteins are shown in blue (adjusted *p*-value < 0.1, log_2_ fold change < -1). The rest of the proteins are shown in light gray. **c,** Gene Ontology (GO) analysis of the differentially enriched proteins in R475G condensates, illustrated as orange dots in **b**. **d,** Gene Ontology (GO) analysis of the differentially depleted proteins in R475G condensates, illustrated as blue dots in **b**. **e,** Representative images of localization of mCherry-tagged TRIM32 and ANKRN52 in mClover3-tagged WT- or R475G-expressing HeLa cells. Scale bar, 10 μm.

APEX2-WT DDX3X and APEX2-R475G were expressed in cells at a similar level (**Extended Data Fig. 5a**). After incubating with biotin-phenol for 30 min, cells were exposed to H_2_O_2_ for 1 min to trigger biotin labelling by the APEX2 moiety. Cells were then lysed, and the labeled proteins were enriched via streptavidin pulldown. Protein labeling and enrichment were analyzed by western blots before being subjected to proteomics analysis (**Extended Data Fig. 5b**). The Western blots and Coomassie stain results showed that the biotinylated proteins were enriched in pulldown fractions compared to the flowthrough and the input samples (**Extended Data Fig. 5b**).

Unsupervised hierarchical clustering of the APEX2-MS protein abundances resulted in two separate clusters for the biological triplicates of APEX2-WT and APEX2-R475G, indicating that R475G mutation was the main determinant of differences between the interactomes (**Extended Data Fig. 5c**). Proteins with significantly altered abundance were selected based on the following criteria: (1) minimum fold-change of 2 in either direction, adjusted *p*-value < 0.1, (2) identified by a minimum of 2 peptides, and (3) detected in at least two replicates (**Supplementary Table 1**).

Analysis of these significant proteins revealed that, compared to the APEX2-WT DDX3X pulldown, 33 proteins were enriched in APEX2-R475G hollow condensates while 128 proteins were enriched in the APEX2-WT pulldown (**Fig. 5b**). Gene ontology (GO) analysis suggests that the proteins enriched in R475G condensates participate in the regulation of Rac protein signal transduction and protein polymerization, while proteins depleted in R475G condensates mainly play roles in translation regulation and ribonucleoprotein complex biogenesis (**Fig. 5c and 5d**).

Because these hollow condensates appear biphasic, with the potential for proteins to either accumulate in the hollow core or the spherical shell of the puncta, we next sought to determine where in the condensates the proteins identified in our proteomics screen might reside. In a previous study of TDP-43 hollow condensates, it was determined that conventional antibody-based immunofluorescence cannot be used to detect the components in the hollow core, as fixation produces a barrier that prevents antibody penetration^42^. Thus, TRIM32 (an E3 ubiquitin ligase) and ANKRN52 (a regulatory protein of protein phosphatase 6), two top targets enriched in APEX2-R475G mass spec pulldown (**Fig. 5b**), were constructed with an mCherry tag, and then subjected to fluorescence imaging. In WT DDX3X-expressing cells, TRIM32 and ANKRN52 were mostly diffuse in the cytoplasm, with TRIM32 showing small foci. In contrast, in cells expressing R475G, TRIM32 and ANKRD52 were sequestered inside of R475G hollow condensates (**Fig. 5e**). The trapping of these proteins in the relatively undynamic R475G condensates may disrupt the cellular pathways in which they normally participate.

### DDX3X mutant hollow condensates recruit WT DDX3X and DDX3Y

As the vast majority of DDX3X disease mutants are expressed heterozygously^4,43^, we wondered whether DDX3X disease-related mutants co-condense with WT DDX3X or DDX3Y in cells. To assess this, we first co-expressed mCherry-tagged WT DDX3X with each of the mClover3-tagged hollow mutants at a similar level in HeLa cells to look for the presence of co-condensates. When co-expressed, WT DDX3X was recruited to the shells of all the hollow condensates formed by each mutant (**Fig. 6a**). The presence of these co-condensates without addition of a stressor suggests that these DDX3X mutants could contribute to disease (at least partially) by trapping WT DDX3X within their hollow condensates. To assess the ability of WT DDX3X molecules to move between the hollow condensates and the cytoplasm, we measured the dynamics of WT DDX3X sequestered in the hollow condensates using full bleach FRAP experiments. WT DDX3X recovered to ∼80% of its original intensity when in co-condensates with G302V, G325E, A497V, and R534H. This is comparable to the level of FRAP recovery of DDX3X in SGs^19^. However, WT DDX3X only recovered to ∼40% of its original intensity when in co-condensates with R475G (**Extended Data Fig. 6a and Fig. 6c**, **left**). While all the hollow mutant condensates interfere with the cellular localization of WT DDX3X, condensates of R475G appeared to trap WT more strongly than other mutants.

**Fig. 6:**
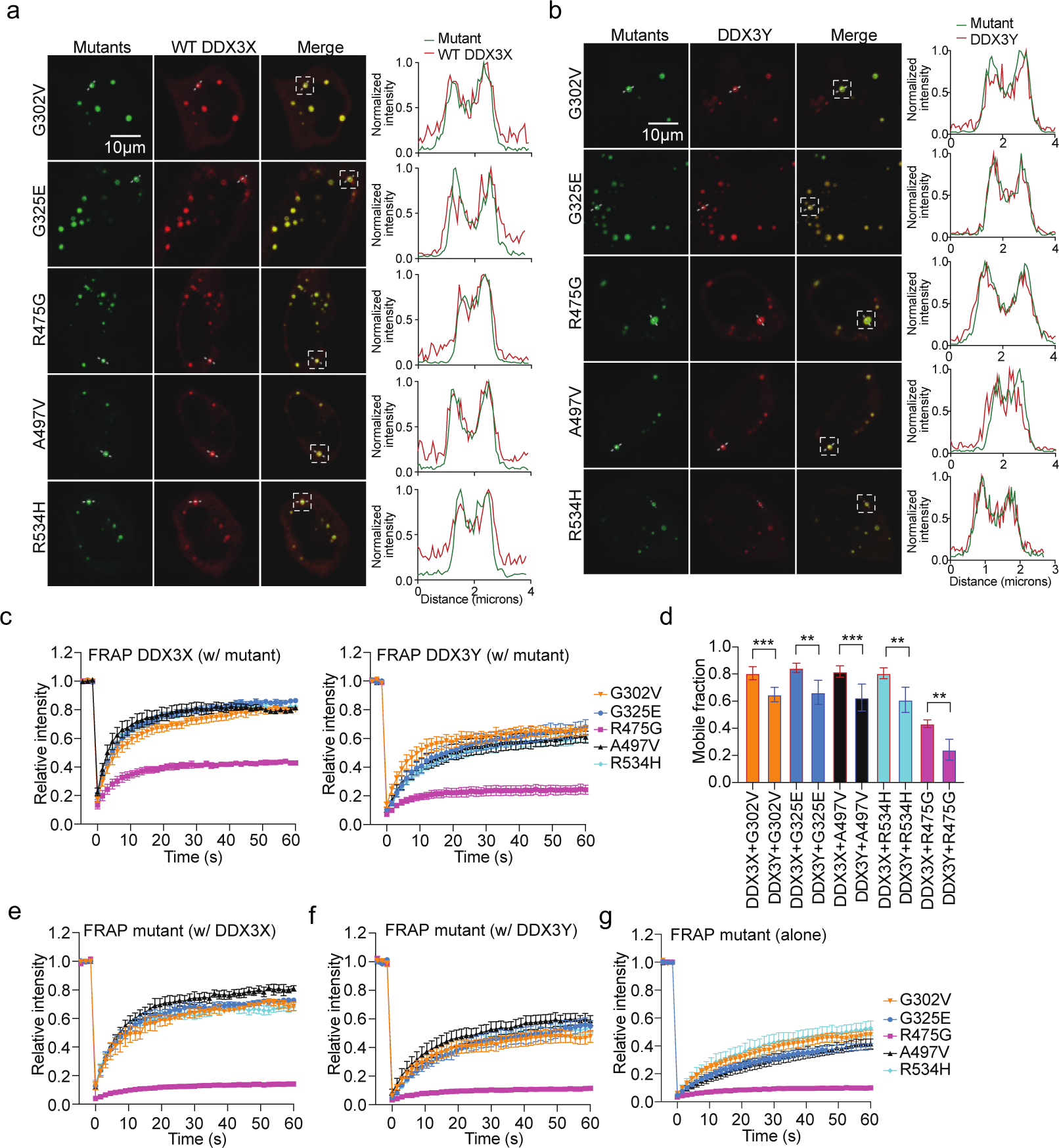
Mutant hollow condensates trap WT DDX3X and DDX3Y. **a,** Representative images showing co-localization of mCherry-tagged WT DDX3X and mClover3-tagged DDX3X hollow variants. White boxes indicate the condensates used to generate the corresponding intensity plots. The intensity plots were generated along the dotted line in the white box. Scale bar, 10 μm. **b,** Representative images showing co-localization of mCherry-tagged DDX3Y and mClover3-tagged DDX3X hollow variants. White boxes indicate the condensates used to generate the corresponding intensity plots. The intensity plots were generated along the dotted line in the white box. Scale bar, 10 μm **c,** Recovery curves for WT DDX3X (left panel) and DDX3Y (right panel) sequestered in the hollow condensates formed by mClover3-tagged DDX3X hollow variants in HeLa cells. The traces of the FRAP data represent means ± s.e.m. from at least three biologically independent experiments. **d,** Comparison of the mobile fractions of mCherry-tagged WT DDX3X or DDX3Y in condensates with mClover3-tagged DDX3X variants, taken from the FRAP analysis in c. The mobile fraction was calculated as described in the methods. Values represent means ± from at least three biologically independent experiments. Significance was calculated using a two-tailed t-test. \*\**p*<0.01; \*\*\**p*<0.001. **e,** Recovery curves for each mClover3-tagged DDX3X variant in hollow co-condensates with mCherry-tagged WT DDX3X in HeLa cells from the cellular FRAP experiments. The traces of the FRAP data represent means ± s.e.m. from at least three biologically independent experiments. **f,** Recovery curves for each mClover3-tagged DDX3X variant in hollow co-condensates with mCherry-tagged DDX3Y in HeLa cells from the cellular FRAP experiments. The traces of the FRAP data represent means ± s.e.m. from at least three biologically independent experiments. **g,** Recovery curves for each mClover3-tagged DDX3X variant in condensates when expressed alone in HeLa cells from the cellular FRAP experiments. The traces of the FRAP data represent means ± s.e.m. from at least three biologically independent experiments.

We next performed these experiments with DDX3Y. Co-expression of mCherry-tagged DDX3Y and each of the mClover3-tagged DDX3X mutants showed that, as with WT DDX3X, DDX3Y was recruited to the shells of each mutant’s hollow condensate (**Fig. 6b**). In our previous work, we found that condensates of DDX3Y (both *in vitro* and in cells) were less dynamic than condensates of DDX3X^19^. We wondered whether the same would be true for DDX3Y in co-condensates with DDX3X mutants. Thus, we repeated the full bleach FRAP in HeLa cells co-expressing DDX3Y and each of the DDX3X mutants. (**Extended Data Fig. 6b and Fig. 6c**, **right**). The results with DDX3Y largely mirrored those with WT DDX3X: the sequestered DDX3Y in the G302V, G325E, A497V, and R534H condensates was more dynamic than in R475G condensates (**Fig. 6c, right**). However, WT DDX3Y only recovered to ∼60% of its original intensity when sequestered in G302V, G325E, A497V, and R534H hollow condensates, and ∼20% when sequestered into R475G condensates. The mobile fraction of DDX3Y was significantly lower than the mobile fraction of WT DDX3X in each type of co-condensate measured (**Fig. 6d**), although there was no significant difference in recovery halftime for G302V, A497V, and R475G mutants when they co-condense with either DDX3X or DDX3Y (**Extended Data Fig. 6c**). The reduced dynamics of these co-condensates suggests that sequestration of WT DDX3X or DDX3Y within hollow condensates may inhibit their ability to facilitate translation, as both WT DDX3X and DDX3Y participate in translation initiation^44^.

Reciprocally, we performed full bleach FRAP on DDX3X hollow mutants in co-condensates with WT DDX3X or DDX3Y. When co-expressed with mCherry-tagged WT DDX3X (**Extended Data Fig. 6d and Fig. 6E**), all the hollow mutants showed improved mobile fractions compared to when they were expressed alone (**Extended Data Fig. 6f**, **Fig. 6g, and Extended Data Fig. 6g**), although the magnitude of this effect was much smaller for R475G than it was for the other hollow mutants. However, when co-expressed with mCherry-tagged DDX3Y (**Extended Data Fig. 6e and Fig. 6f**), only G325E showed significant improvement in its mobile fraction (**Extended Data Fig. 6g**). Additionally, both DDX3X and DDX3Y were able to reduce the recovery halftime for all the hollow mutants except for R475G (**Extended Data Fig. 6h**). These data indicate that co-condensates of WT DDX3X and mutant DDX3X (which would be present in XX individuals with a mutated *DDX3X* allele) are more dynamic than co-condensates of DDX3Y and mutant DDX3X (which would be present in XY individuals with a mutated *DDX3X* allele).

### Hollow mutations of DDX3X are over-represented in cases of DDX3X-related cancers and neurodevelopmental disorders

Given the striking morphological and catalytic phenotypes observed in our hollow mutants, we questioned whether these mutants may be correlated more strongly with DDX3X-releated diseases than the diffuse mutants. We examined the cBio Portal database^45^ along with several studies of DDX3X-related neurodevelopmental disorder (NDD)^4,26,46,47^ and counted the number of patient samples in which mutations at the position of each hollow mutant appeared. Across this data set, mutations at the positions of the hollow mutants accounted for 60% of total appearances, while mutations at the positions of the diffuse and speckled positions represented only 31% and 9% of appearances, respectively (**Fig. 7a**). If the three morphologies were all equally likely to appear in disease, one would expect hollow mutants to appear ∼42% of appearances (as there are five hollow mutants out of the twelve investigated in this study), indicating that hollow mutants are over-represented in, and likely more causative of, disease. Thus, DDX3X mutations which impair catalytic activity and cause aberrant cellular phase separation are possibly a feature of DDX3X-related diseases.

**Fig. 7:**
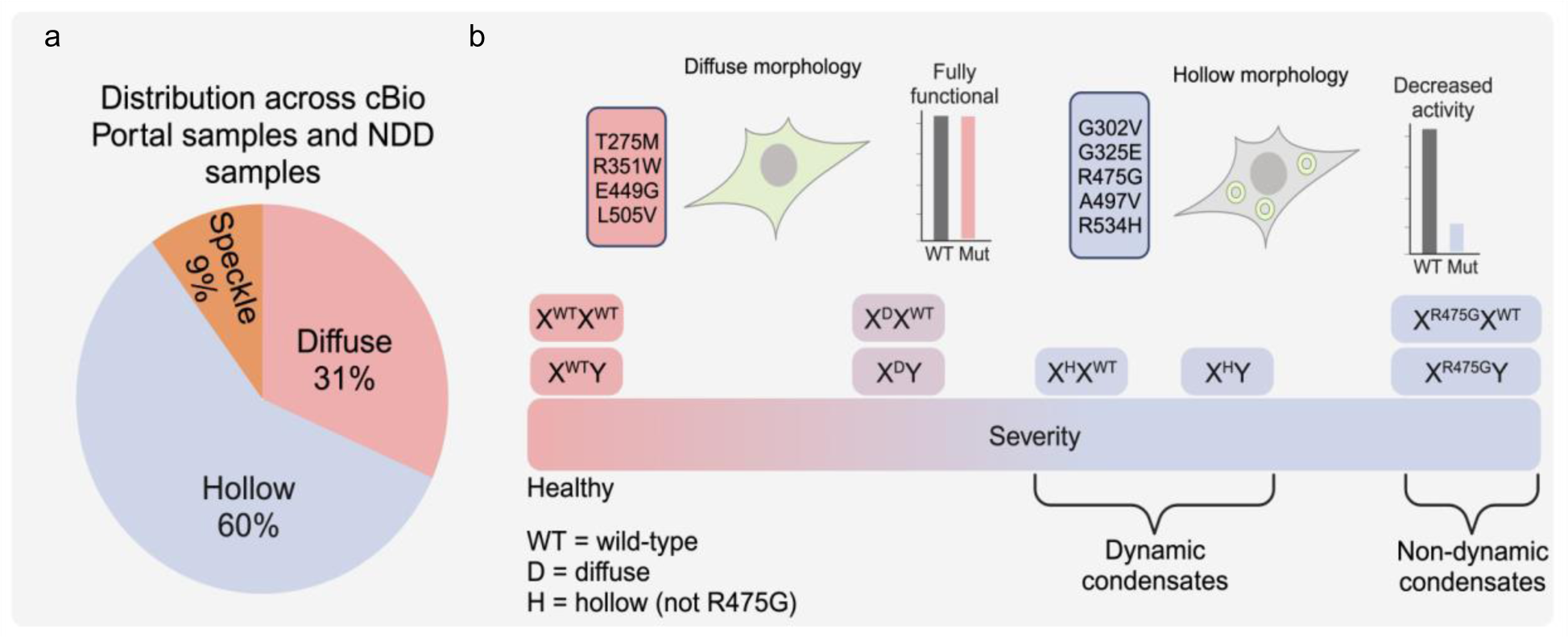
Distribution of mutants across DDX3X-related diseases and proposed model. **a,** Distribution of mutants in this study across cBio Portal samples (cancer) and neurodevelopmental studies. **b,** Proposed model. DDX3X mutants that disrupt ATPase and strand release activities form hollow condensates in cells without external stress, while mutants that do not disrupt catalysis remain diffuse. The severity of disease related to these mutations is correlated with the dynamics of these condensates. When coexpressed with WT DDX3X, hollow puncta are more dynamic than when coexpressed with DDX3Y, which may explain sex bias in DDX3X-related diseases.

## Discussion

Our work introduces an integrative approach across structural, biochemical, biophysical, and cellular methods to categorize DDX3X disease mutants. In doing so, we found that a specific group of DDX3X mutants that are over-represented in human diseases form cellular hollow condensates. We further revealed the molecular mechanism underlying the formation of these condensates: deficient ATPase and RNA strand release activities relative to wild-type protein. Hollow condensates sequestered WT DDX3X, DDX3Y, and proteins in diverse cell signaling pathways, possibly providing a reason why these mutants are over-represented among diseases associated with DDX3X mutations. Furthermore, we found that the presence of WT DDX3X improved the dynamics of heterogenous hollow condensates ∼20% more than the presence of DDX3Y, underscoring that differential interactions with WT DDX3X and DDX3Y may underlie sex biases in DDX3X-related disease (**Fig. 7b**).

### Select DDX3X mutants form unique cytoplasmic hollow condensates in cells

The formation of membraneless compartments within cells via liquid-liquid phase separation is crucial for the proper regulation of several cellular processes, ranging from gene expression to signal transduction^48,49^. Some condensates, such as SGs, have a uniform spherical morphology in which each component is fairly well-distributed^50^, while others have distinct sub-compartments, such as the nucleolus^51^. In addition, a different class of multilayered condensates has been reported where the condensate forms a vesicle-like structure with a seemingly lower content (“hollow”) internal space^37,42,52^. Until this point, these hollow cellular condensates have only been observed in the nucleus when seen in cells^42^. In our current study, we found that a specific subset of disease mutants of DDX3X can form hollow condensates in the cytoplasm and these condensates may disturb function of WT DDX3X, DDX3Y, and protein signal transduction (**Figs. 5 and 6**).

Formation of these hollow condensates can be either a passive process regulated by the protein’s own biochemical and/or biophysical properties or an active process driven by “chaperone” proteins^42^. In the case of *in vitro* condensates of protamine, hollow condensates are formed via the liquid-liquid phase separation of protein-RNA complexes when the RNA:protein ratio is either >1.87 or <0.075^52^. On the other hand, RNA binding-deficient mutants of TDP-43 require the ATPase activity of HSP70 to form hollow condensates^42^. For the DDX3X mutants tested in this study, we found that the propensity to form hollow condensates was correlated with the proteins’ decreased ATPase and/or strand release activities (**Fig. 2**), and that the dynamics of these hollow condensates may be linked to their RNA binding affinities (**Fig. 4**). We did not find any chaperone-like proteins inside of the hollow condensate in our APEX2-MS analysis (**Fig. 5**). However, DDX3X is an ATPase, unlike TDP-43. It is possible that mutants with diminished strand release activity form hollow condensates because they are unable to release bound RNA. In the case of R475G, its starkly diminished RNA binding capacity (**Figs. 2e and 4d**) results in condensates with less unreleased RNA than condensates of other hollow mutants. Although it has been previously shown that increased RNA concentrations in condensates of the DEAD-box helicase Dhh1 resulted in decreased dynamics^53^, this is not the case for R475G, which has the lowest dynamics of all mutants tested. It is possible that the relationship between internal RNA concentration and condensate dynamics is different for human DDX3X and its mutants than it is for yeast Dhh1. This difference may be driven by their intrinsically disordered regions, as we have previously should that differences in IDR sequences are responsible for the different dynamics of DDX3X and DDX3Y condensates^19^. Because our hollow mutants are over-represented in disease (**Fig. 7**), revealing the cause of their condensation may provide important insights for future therapeutic development.

### Hollow mutations of DDX3X cause disruptions throughout its catalytic cycle

To date, several crystal structures of DDX3X (and other closely related DEAD-box proteins) with different combinations of ATP analogs and RNA substrates have been deposited to the PDB. Using these structures, a model of DDX3X’s catalytic cycle was recently described in which DDX3X binds dsRNA as a dimer, then performs strand separation upon subsequent binding of ATP. The post-unwound ATP:DDX3X:ssRNA ternary complex is then disassociated by the hydrolysis of ATP and the release of ssRNA and ADP (**Fig. 2b**). Based on this reaction scheme, we were able to directly assess how our different DDX3X mutations impair different parts of the catalytic cycle. In particular, we found that R475G disrupts the transition from the apo state to the pre-unwound state (**Fig. 2b, step 1**) by disrupting RNA binding in addition to other potential effects (**Fig. 3b**), G325E disrupts the transition from the pre-unwound state to the post-unwound state (**Fig. 2b, step 2**) by blocking the deposition of ssRNA in the acceptor cleft (**Fig. 3e**), and R534H explicitly blocks the hydrolysis of ATP (**Fig. 2e**, **step 3**) as it does not cause defects in RNA binding (**Fig. 2e**) or ATP binding (**Fig. 3h)**. All of these defects stall the catalytic cycle and limit ATP hydrolysis and strand release (**Fig. 2c and 2f**). Many of these mutations are also linked with disease severity. R475G has been noted to cause severe neurodevelopmental defects, as has been noted with mutations in close proximity to G325E (T323I and R326H) and R534H (T532M)^4^. Collectively, these results provide evidence that disruptions in DDX3X’s catalytic cycle and hollow condensate formation are linked to disease severity.

Beyond the implications for disease, these findings shed light on the role of these residues in the catalytic cycle of DDX3 (and DEAD-box proteins in general, as these amino acids are largely conserved). These disease-related mutants provided us with natural experiments to test both the role of particular DEAD-box residues and the current model of DDX3-mediated catalysis as a whole (**Fig. 2b**). Taken together with our concurrent in-depth investigation of how the intrinsically disordered regions of DDX3X and DDX3Y both contribute to enzyme catalysis and differentiate the activities of these sexually dimorphic homologs via the formation of newly-described protein-RNA clusters (RPCs)^20^, our work provides the most complete picture to date of how the structured and intrinsically disordered regions of DDX3X and DDX3Y contribute to enzyme activity.

### Hollow condensates of DDX3X mutants may contribute to disease via sequestration of other proteins

Under unstressed conditions, WT DDX3X is diffuse in cells, whereas upon addition of stressors such as oxidative stress or energy depletion, it is compartmented into G3BP1-positive SGs^7^. In contrast, the five hollow mutants identified herein formed their cellular condensates without addition of an external stressor. These hollow condensates do not contain G3BP1, meaning that they are not “canonical” SGs^50^. This raised the question as to what role these hollow condensates play in disease. To address this question, we performed APEX2 proximity labeling followed by mass spectrometry to identify the other protein components of hollow condensates formed by DDX3X R475G (**Fig. 5**). The results revealed that R475G hollow condensates sequester a subset of proteins that do not normally interact with WT DDX3X. R475G condensates are also depleted of proteins relating to protein translation, further separating them from SGs^50^.

One of the most enriched proteins in R475G condensates was the E3 ubiquitin ligase TRIM32. During neuronal development, TRIM32 drives neural progenitor differentiation by preferentially accumulating in one daughter cell during division. The TRIM32-positive daughter cell will differentiate into a mature neuron, while the TRIM32-negative daughter cell will remain a progenitor^54^. Expression of R475G in neural progenitor cells might disrupt the proper segregation of TRIM32, as its hollow condensates are significantly less dynamic than condensates of other mutants (**Figs. 4 and 6**). This possibly contributes to R475G’s more severe neurodevelopmental phenotypes than some other DDX3X mutants. In addition to TRIM32, many of the other proteins enriched in R475G condensates are involved in the regulation of Rac signaling. DDX3X mutants may thus also dysregulate Rac signaling, which has been implicated in a number of cancers^55^. The disease implications of these sequestered proteins warrant further investigation.

### The distinct properties of DDX3X mutant hollow condensates may correlate with sex biases in disease incidence and severity

Beyond the protein components identified in our APEX2-MS screen, fluorescence imaging revealed that DDX3X mutant hollow condensates sequester WT DDX3X or DDX3Y when co-expressed in cells (**Fig. 6**). FRAP experiments further revealed that co-condensates of mutant DDX3X and WT DDX3X are more dynamic than co-condensates of mutant DDX3X and DDX3Y (**Fig. 6**). These differing condensation properties may contribute to the sex biases in incidence and severity of DDX3X-related disorders, which are frequently biased against XY-individuals^4,21^. The disruption of cellular condensate dynamics can often lead to tumorigenesis, as is the case with mutant forms of UTX^23^ and AKAP95^56^. Additionally, a mutant of DDX3X not included in this study (L566S) was recently found to form cellular condensates with dynamics comparable to R475G and to form amyloid-like fibrils *in vitro*^57^, further supporting a link between DDX3X condensation and disease. Based on our results, we hypothesize that DDX3X mutant hollow condensates formed in XY individuals would be less dynamic and more able to sequester the remaining DDX3Y (and other constituent proteins) than condensates formed in XX individuals. As such, we propose that mutants of DDX3X that form less dynamic condensates are more likely to contribute to severe disease, and that this effect is exacerbated by co-expression with DDX3Y, as in any person with a Y chromosome (**Fig. 7b**). Future work should prioritize the study of these mutants in both XX and XY chromosomal sex-defined genetic backgrounds. Collectively, the results of this study indicate that the enzymatic properties of select DDX3X disease mutants contribute to the formation of cytoplasmic hollow condensates with unique protein components which may contribute to sex biases in human disorders.

## AUTHOR CONTRIBUTIONS

M.C.O., H. Shen, and K.F.L. designed the experiments. H. Shen, A.Y., M.C.O., and E.L. generated the constructs and the mammalian cell lines used in this study. H. Shen, A.Y., M.C.O., E.L., and X.W. purified the recombinant DDX3X and DDX3X mutants used in this study. H.S. and A.Y. performed the immunofluorescence imaging and FRAP experiments and analyzed the data. M.C.O. performed the ATP binding assay, the ATPase assays, and the EMSA assays together with E.L.. M.C.O. performed the structural analysis and generated Supplemental Movie 1. M.S.M.F. and M.C.O. performed the plasmid titration of Western blots and imaging in Fig. 1. A.Y. performed the fluorescence anisotropy experiments with guidance from Y.E.G. and H. Shweta. H. Shen performed the APEX2 proteomics and H.Y.T performed the proteomic data analysis. All authors participated in writing, discussing, and editing the manuscript.

## DATA AVAILABILITY

Any data supporting the findings of this study and additional information required to reanalyze the data reported in this paper is available from the lead contact upon request.

## ACKNOWLEDGEMENTS

This work was supported by the National Institutes of Health (R35GM133721 and R01HL160726 to K.F.L., R35GM133721-03S1 to A.Y., R50CA221838 to H.Y.T, T32GM132039 to A.Y. and M.C.O., and R35GM118139 to Y.E.G.). K.F.L. is supported by the American Cancer Society (RSG-22-064-01-RMC), the Damon Runyon Innovator Award, and the Linda Pechenik Montague Investigator Award. We thank Dr. Matthew Kayser for sharing the Leica SP8 confocal microscope. We would also like to thank Drs. Roger Greenberg, Ben Black, and Nancy Bonini for their helpful comments while drafting this manuscript. Fig. 1a and Fig. 7 were created with Biorender.com.

## METHODS

### Cell culture, transfection, and *Escherichia coli* strains

HeLa and HEK293T cells were cultured in DMEM + GlutaMAX (GIBCO) with 10% FBS (GIBCO) and 1% Pen/Strep (Corning) in a humidified incubator with 5% CO_2_ at 37°C. Lipofectamine 2000 (Invitrogen) was used for plasmids transfection. Experiments were conducted on transfected cells after 24 hrs transfection. For bacterial cell culture, TOP10 and BL21(DE3)-RIL chemically competent bacterial strains were grown in lysogeny broth containing the corresponding antibiotics at 200 rpm, 37°C.

### Constructs

The DDX3X-mCherry plasmid was generated in our previous study (Shen et al., 2022) and the DDX3X-mClover3 was constructed by substitute the fluorescent tag using fusion PCR. The different mClover3-tagged DDX3X mutants were generated using fusion PCR with the wild-type DDX3X-mClover3 plasmid as template. For the APEX2-MS experiments, the DNA fragment encoding APEX2 was PCR amplified from pcDNA5/FRT/TO APEX2-GFP (Addgene) and fused to the N-terminus of DDX3X using fusion PCR. To verify the specificity of APEX2 labeling, TRIM32 and ANKRD52 genes were cloned from HeLa cDNA, fused with a mCherry tag at the C terminus, and inserted into the pcDNA3 vector for mammalian cell expression. G3BP1 was also cloned from HeLa cDNA, fused with an mCherry tag at the C-terminus, and inserted into the pPB vector. All plasmids were validated by Sanger sequencing.

### Protein purification

Wild-type and mutant DDX3X tagged with mCherry were purified according to our previous report (Shen et al., 2022). Briefly, the plasmids were transformed into *Escherichia coli* strain BL21-RIL to express the recombinant proteins. The bacteria were cultured in lysogeny broth at 37°C until OD_600 nm_ = 0.8 before administration of 1 mM IPTG and shaken at 200 rpm at 16°C for 16 hrs after induction. The pellets from 2 L bacterial culture were resuspended with 80 mL binding buffer (25 mM Tris-HCl, pH 7.5, 10% glycerol, 500 mM NaCl) and subjected to sonication. After centrifuging at 12,000 rpm for 30 min to remove the cell debris, the supernatant was loaded onto a Ni-NTA column. Next, 10 column volumes of the binding buffer supplemented with 50 mM imidazole was used to wash away the non-specifically bound proteins. Another 10 column volumes of a high salt buffer (25 mM Tris-HCl, pH 7.5, 10% glycerol, and 2 M NaCl) was used to remove the associated RNAs from wild-type DDX3X and DDX3X mutants. Finally, four column volumes of the binding buffer supplemented with 500 mM imidazole was used to elute the bound proteins, which were dialyzed into the storage buffer (25 mM Tris-HCl, pH 7.5, 500 mM NaCl, 10% glycerol, and 2 mM DTT) with TEV enzyme cleavage of the His-tag simultaneously for 1 hr at room temperature. Then, the proteins were concentrated using Amicon Ultra-15 (Millipore) tubes before loading to a Superdex 75 column, with storage buffer (25 mM Tris-HCl, pH 7.5, 500 mM NaCl, 10% glycerol, and 2 mM DTT). The purity of the proteins was analyzed by SDS-PAGE. Purified proteins were aliquoted, snap-frozen in liquid nitrogen, and stored at -80 °C. Once thawed, aliquots were never refrozen. All purification steps after sonication were performed at room temperature.

### Electrophoretic mobility shift assay (EMSA)

To measure the RNA binding affinities of different DDX3X mutants, EMSAs were carried out. A double-stranded RNA probe with 5’ overhang, to which DDX3X is known to bind^19,58^, with following sequences: 5’-biotin/ACCGCUGCCGUCGCUCCG/AlexF647N/-3’ and 5’-/Cy3/UUUUUUUUUUUUUUUUUUUUUUUUUUUUUUUUUUUUUUUUUUUUUUUUUUCGGAGCGACGGCAGCGGU-3’ was used. Both strands were ordered from IDT and annealed by heating the equimolar mixtures of both strands at 65°C for 5 min, then slowly cooling the sample to room temperature over 3 hrs. Various concentrations of each mCherry-tagged DDX3X mutant proteins were incubated with 100 nM double-stranded RNA probe at room for 30 min in a buffer consisting of 50 mM Tris-HCl pH 7.5, 150 mM KCl, 2 mM MgCl_2_, 10 mM beta-mercaptoethanol and .1 mg/mL BSA before loading to a homemade 6% polyacrylamide gel (37.5:1) in 0.5 × TBE buffer. The gel was run at 120 V for 2 hrs at 4°C. The gel was then scanned using an Amersham Typhoon using the Cy5 channel. Fiji was used to quantify intensity of the bottom free RNA band. Fraction bound was calculated using the following equation: Fraction bound = 100*(1-(I_n_/I_o_)), where “I_n_” is the intensity of the free probe band at a given concentration and “I_o_” is the intensity of the free probe band at 0 nM protein. Fraction bound was plotted versus concentration in GraphPad Prism and fit with the Hill equation to give *K_D_*.

### Fluorescence imaging

HeLa cells were seeded to a 6-well plate with a coverslip in each well and cultured overnight. The various DDX3X plasmids were transfected to cells with Lipofectamine 2000 the next morning. After 24 hrs in culture, the cells were washed once with PBS and then fixed using 4% paraformaldehyde in PBST (PBS with 0.01% Tween-20) at room temperature for 15 min. Then, the cells were washed twice by PBST and permeabilized by 0.5% Triton in PBST at room temperature for 20 min. After being washed once with PBST, the cells were blocked with 1% BSA in PBST at room temperature for 30 min. After being washed three times with PBST, the cells were incubated with 0.5 μg/mL DAPI for 1 min. After 4 PBST washes, an antifade reagent (Invitrogen) was used to mount the slides. The images were taken using a Leica TCS SP8 confocal microscope.

### Quantification of hollow puncta

To quantify the percentage of cells showing hollow puncta, a slide with HeLa cells transiently transfected with each DDX3X mutant was scanned left to right and top to bottom. The images for the first 25 transfected cells with puncta were taken. We then calculated the number of cells with hollow puncta divided by the total 25 cells. For each DDX3X mutant, three biological replicates were carried out. The results were plotted using GraphPad Prism.

### EU labeling of RNA in cells

To quantify the RNAs in DDX3X mutant condensates, the cellular RNAs were first labeled with 5-ethynyl uridine (EU). An Alexa Fluor 594 fluorophore was then added through a click chemistry reaction using Click-iT RNA image kits (ThermoFisher). Briefly, HeLa cells were seeded to a 6-well plate. The plasmids of DDX3X mutants were transfected into the cells using Lipofectamine 2000. After 24 hrs in culture, the medium was removed and fresh medium containing 1 mM EU was added. After a 1 hr incubation, the cells were fixed and permeabilized as stated above. Alexa 594 was conjugated to EU using the Click-iT reaction. After three PBST washes, the cells were incubated with 1 mL Hoechst 33342 to stain the DNA. After 4 PBST washes, an antifade reagent (Invitrogen) was used to mount the slides. The images were taken using a Leica TCS SP8 confocal microscope. The average Alexa 594 intensity in hollow puncta of each DDX3X mutant was quantified using Fiji ^59^, with the average Alexa 594 intensity in the corresponding nucleoplasm used for normalization.

### Fluorescence recovery after photobleaching (FRAP)

The FRAP assays were conducted using the bleaching module of a Zeiss LSM 880 confocal microscope. The 488 nm laser was used to bleach the mClover3 signal, and the 561 nm laser was used to bleach the mCherry signal. HeLa cells were seeded on 35 mm poly-D-lysine coated glass-bottomed dishes (Mattek), and the plasmids of DDX3X mutants were transfected into the HeLa cells. After 24 hrs culture, the normal DMEM medium was replaced with FluoroBrite DMEM (GIBCO) medium with 10% FBS. For full bleaching assay to dissect the molecule exchange between puncta and surroundings, an entire punctum was selected to bleach with 100% laser power and time-lapse images were collected afterward. To probe the molecule exchange inside of the puncta, partial bleaching was performed by focusing on a circular region of interest (ROI) of a punctum using 100% laser power, and time-lapse images were collected afterward as well. The fluorescence intensity was directly measured in the Zen software. The values were reported relative to pre-bleaching time points. GraphPad Prism was used to plot the data. The halftime for each replicate was calculated using the following formula: y=a•(1-exp(-b•x)) + c, in which “a” is the slow recovery fraction, “c” is the rapid diffusion fraction, and “b” is the recovery rate. The halftime is ln2 / b, and the mobile fraction is a + c. Significance was calculated using an unpaired Student’s two-tailed t test.

### Malachite green ATPase assay

ATPase measurements were taken using the Malachite Green Phosphate Assay Kit (Sigma) according to the manufacturer’s instructions. Briefly, 1 μM of each mCherry tagged DDX3X mutant protein was incubated with 100 ng/μL total RNA extracted from HeLa cells for 15 min in the reaction buffer (25 mM Tris-HCl, pH 8, 200 mM NaCl, 1 mM DTT, and 2 mM MgCl_2_) before the addition of 2 mM ATP. The reaction was incubated at 16°C for 30 min. Then the reaction was quenched by the addition of malachite green mixture and left for an additional 30 min to develop the color. The samples were then loaded into a clear-bottom 384-well plate and the absorbance at OD_620_ nm was measured. Values were converted from absorbance units to μM free phosphate using a standard curve generated with the kit’s phosphate standard. Background values (free phosphate detected from reactions lacking RNA) were subtracted from the values from reactions with RNA. Data were plotted in GraphPad Prism. Significance was calculated using an unpaired Student’s two-tailed t test.

### Multi-parameter confocal fluorescence time-resolved microscopy and spectroscopy

All single molecule measurements were performed on multi-parameter confocal time-correlated single photon counting (TCSPC) microscopy and spectroscopy (MicroTime-200; PicoQuant, GmbH). Double stranded RNA: an 18-mer (5’-biotin/ACCGCUGCCGUCGCUCCG/Alexa647N/-3’) annealed to a 42-mer (5’-/Cy3/UUUUUUUUUUUUUUUUUUUUUUUUCGGAGCGACGGCAGCGGU-3’) (IDT), labelled with the donor (Cy3) and acceptor (Alexa647) dyes were excited with 532 nm and 637 nm pulsed diode lasers (LDH-D-TA-532, LDH-D-TA-637, PicoQuant) in pulsed-interleaved excitation (PIE) mode at a repetition rate of 20 MHz. The beam passes through an excitation dichroic filter ZT532/637 (Chroma Technology) and a water immersion objective lens Olympus UPLanSApo 60x/1.2 w with collar correction for coverslip thickness. All the measurements were carried out with laser focused 20 μm above the coverslip, while maintaining the power of both the lasers below 20 μW. Fluorescence signals from the sample after passing through 50 μm pinhole were split into vertical and horizontal channels by polarized beam splitter (U-MBF3-Olympus). Cy3 and Alexa647 emission signals were separated by identical dichroic filters (T635 lpxr, Chroma Technology) in each pathway. Polarized and wavelength-selected photons through bandpass filters ET582/64 (for donor, Cy3) and ET690/70 (for acceptor, Alexa647) were projected onto four single-photon avalanche photodiode (SPAD) detectors and cataloged by a HydraHarp TCSPC time-interval analyzer in (PicoQuant). Data was acquired for the labeled dsRNA-I alone followed by several 5-minutes recording (over the course of 1 - 2 hrs) after adding the protein to the RNA. Subsequently, data was collected for the course of another hour following the addition of 1 mM MgATP. Nunc Lab-Tek chambers (ThermoFisher-155411) with borosilicate coverslip bottoms were used for all measurements. These chambers were treated with 50% (w/v) PEG-8000 solution, incubated at room temperature for 3 - 4 hours, followed by several washes with FRET buffer (50 mM Tris, pH 7.5, 125 mM NaCl and 2 mM MgCl_2_). Confocal detection volume for 532 and 637 nm lasers were calibrated and determined using rhodamine6-G and atto637N dyes.

### Time-resolved fluorescence anisotropy

The parallel and perpendicular ns fluorescence intensity decays for DO particles, DL particles and AO particles within the respective selected regions of *E vs. S* plots and burst selection filters were used to calculate the time-resolved anisotropy decay (*r*(*t*)) using Equation 9. DO particles used donor excitation (*F_DexDem_*). DL and AO particles used acceptor excitation (*F_AexAem_*).

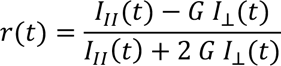

Where, *r* is the anisotropy, *I_II_*(*t*) is the parallel fluorescence intensity, *I_⊥_*(*t*) is the perpendicular fluorescence intensity, *G* is the correction factor for difference in sensitivity of the two detectors. The time resolved anisotropy curves (*r*(*t*)) were fitted with Equation 10 to obtain rotational correlation time (*Φ*) and the proportion molecules with hindered rotation (*B/B_0_*):

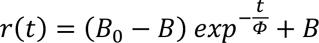

Where, *B*_0_ is the anisotropy value ∼1 ns after the laser pulse and completion internal (segmental) motions within the RNA construct. The value of *B*_0_ was determined from the asymptote (*B*) of recordings with RNA and protein before adding ATP.

Proportion Strand Release (ΔB/B_0_) was calculated by subtracting the end point B/B_0_ (30 minutes after the addition of 1 mM ATP) from the initial B/B_0_ right before 1 mM ATP was added.

### TNP-ATP binding assay

WT DDX3X or R534H (MBP-tagged) was serially diluted to twice the indicated concentrations in 1X reaction buffer (25 mM Tris-HCl, pH 8, 200 mM NaCl, 1 mM DTT, and 2 mM MgCl_2_). TNP-ATP was then diluted to 4 μM in reaction buffer. 50 μL of protein was then mixed with 50 μL TNP-ATP to produce the indicated protein concentration with 2 μM TNP-ATP in 1X buffer. This mixture was then incubated on ice for 10 minutes. The reactions were the loaded into a black bottom 384-well plate and fluorescent emission from 500 – 600 nm was measured with excitation at 410 nm. To analyze data, the emission value at 500 nm was first subtracted from each trace. Then, the value of the no protein sample at 540 nm was subtracted from the other concentration’s values at 540. These values were then plotted vs concentration using GraphPad Prism 9 and fit with a Hill binding curve. Lysozyme, which does not bind ATP, was used as a negative control.

### Cellular APEX labeling

To capture the proteins in the hollow puncta formed by R475G DDX3X, a recently established proximity labeling method was carried out followed by mass spectrometry^19,39^. Briefly, one 10 cm plate of HEK 293T cells was transfected with 3.5 μg APEX2 fused wild-type DDX3X- or R475G- expressing plasmid. After 24 hrs culture, the normal DMEM medium was replaced with DMEM medium supplemented with 500 mM biotin-phenol for 30 min. Then, H_2_O_2_ (Sigma-Aldrich) was added to each cell culture dish at 1 mM final concentration for exactly 1 min with gentle agitation. To stop the labeling, the culture medium was removed, and the quenching solution (10 mM sodium ascorbate, 10 mM sodium azide, and 5 mM Trolox in PBS) was immediately used to wash the cells three times. Finally, 1 mL quenching solution was applied to cover the cells. Cells were then collected with a cell scraper. The unlabeled control samples were prepared in parallel under the same procedure as aforementioned, without the addition of the H_2_O_2_.

### Mass spectrometry sample preparation

The cell pellets collected after APEX labeling were lysed by gentle pipetting in RIPA buffer (50 mM Tris-HCl pH 7.5, 150 mM NaCl, 0.1% (wt/vol) SDS, 0.5% (wt/vol) sodium deoxycholate and 1% (vol/vol) Triton X-100) supplemented with 1x protease inhibitor cocktail, 1 mM PMSF and quenchers (10 mM sodium azide, 10 mM sodium ascorbate and 5 mM Trolox). After resuspension and on ice for ∼ 2 min, the lysates were clarified by centrifuging at 15,000 g for 10 min at 4 °C. The concentration of the cell lysate was quantified by using the Pierce 660-nm assay. The same amount of cell lysate was incubated with RIPA buffer washed Pierce Streptavidin magnetic beads for 1 hr at room temperature on a rotator. The beads were pelleted using a magnetic rack and washed sequentially with 1 mL RIPA buffer twice, once with 1 M KCl, once with 0.1 M Na_2_CO_3_, once with 2 M urea in 10 mM Tris-HCl, pH 8.0, and twice with RIPA lysis buffer. The wash buffers were kept on ice throughout the procedure. The biotinylated proteins were eluted from the beads by boiling each sample in 30 μL of 3 × protein loading buffer supplemented with 2 mM biotin and 20 mM DTT for 10 min. Then, placed the samples on a magnetic rack to pellet the beads and to collect the eluate. All the eluate was loaded to 10% SDS-PAGE gel and ran 1 cm into the gel. Then, the entire stained gel regions were excised, reduced with tris(2-carboxyethyl)phosphine (TCEP), alkylated with iodoacetamide, and digested with trypsin. Tryptic digests were analyzed using a 95 min LC gradient on the Thermo Q Exactive HF mass spectrometer as described previously^60^.

### Mass spectrometry data analysis

MS/MS data were searched with full tryptic specificity against the Swiss Prot human proteome database (06/04/2021), the wild-type DDX3X and R475G protein sequences, and a common contaminant database using MaxQuant 1.6.17.0^61^. Search parameters used include two missed cleavages, static carbamidomethylation of Cys, and variable modifications of protein N-terminal acetylation, Asn deamidation, and Met oxidation. Consensus identification lists were generated with false discovery rates set at 1% for protein, and peptide identifications. High confidence identification of proteins with significant change refers to proteins satisfying the following criteria: minimum fold-change of 2 in either direction, adjusted *p*-value < 0.1, identified by a minimum of 2 razor + unique peptides, and detected in at least two of the three replicates. High confidence proteins were subjected to the gene ontology (GO) analysis using the Metascape^62^.

### Protein quantification and Western blot

Protein concentrations of the samples were calculated using the Bradford Assay (Thermo Fisher, 23246). Protein samples were boiled at 95°C in Laemmli sample buffer for 10 min. After brief centrifugation, the samples were loaded onto SDS-PAGE gels. After running at 180 V for 1 hr, the gels were transferred to membrane by semi-dry transfer apparatus at 20 V for 50 min. Then, the membrane was blocked with 5% milk or BSA in 1 × PBST for 30 mins at room temperature or 4°C overnight. The membrane was then incubated in 3% milk or BSA in 1 × PBST containing the corresponding primary antibody overnight at 4°C. After washing three times with 1 × PBST, the horseradish peroxidase (HRP)-conjugated secondary antibody (1:20,000) in 1% of milk was applied and incubated at room temperature for 1 hr. After washing three times with PBST, the membranes were visualized using ECL Western Blotting Detection Kit (Thermo Fisher).

## QUANTIFICATION AND STATISTICAL ANALYSIS

Images were analyzed using Fiji. All data are presented as the mean ± standard error of mean (s.e.m.) or standard deviation (s.d.) from the independent determinations. The statistical analyses were performed using the GraphPad Prism (GraphPad Software, Inc,; La Jolla, CA, USA). Differences of means were tested for statistical significance with unpaired two-tailed Student’s t-test. \**p* < 0.05; \*\**p* < 0.01; \*\*\**p* < 0.001; \*\*\*\**p* < 0.0001; n.s. means *p* > 0.05.

## Supplemental Tables

**Supplemental Table 1** Summary of WT-DDX3X and R475G APEX2-MS proteomics results.

## Supplemental Movies

**Movie 1** Animation of DDX3X’s catalytic cycle.

## Extended data figures and figure legends

**Extended Data Fig. 1:**
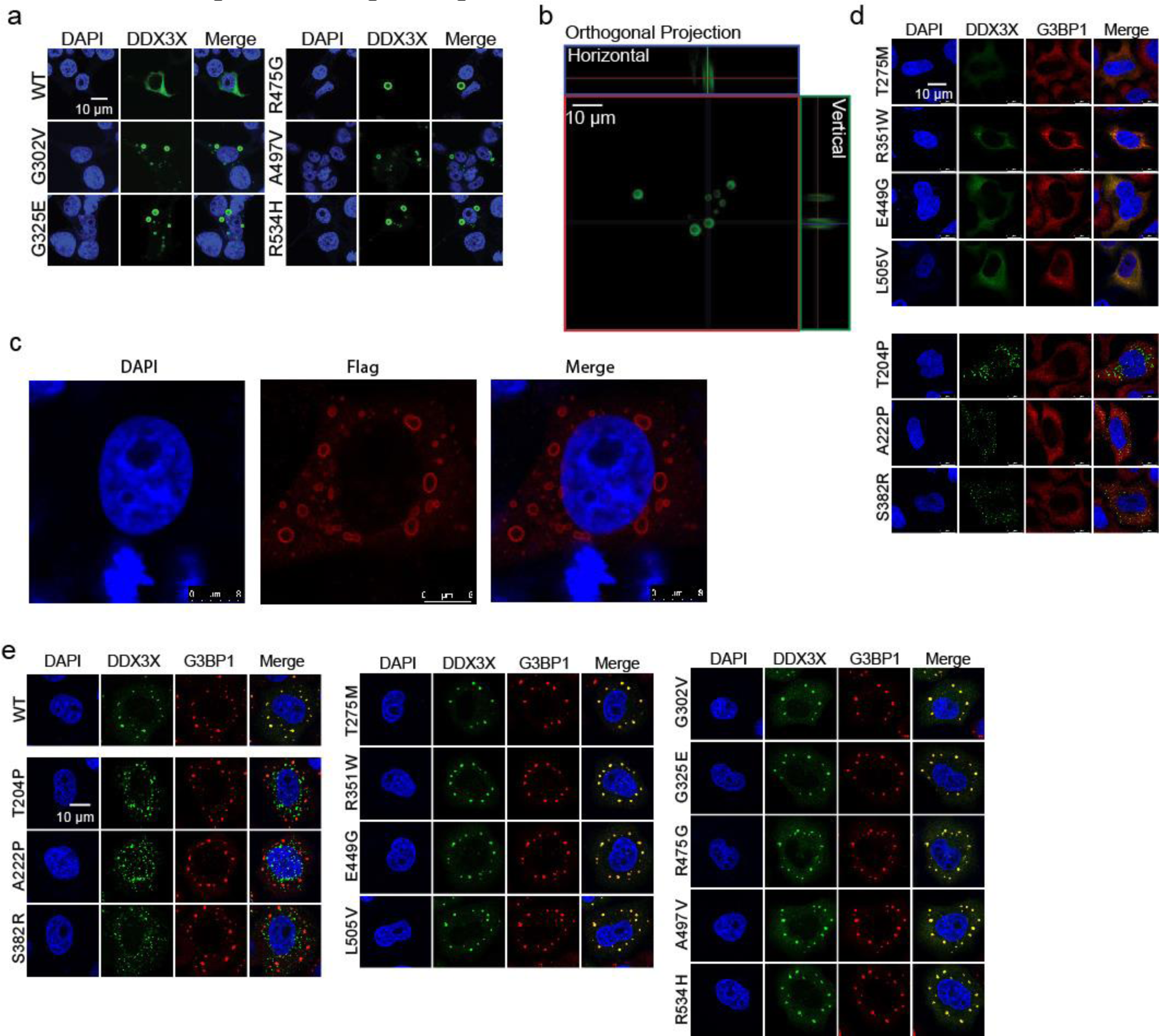
A subset of DDX3X disease mutants form unique hollow condensates in cells. **a,** Representative images of cellular distributions of the mClover3-tagged DDX3X variants in HEK293T cells. Scale bar, 10 µm. **b,** Orthogonal projections demonstrate that mClover3-tagged R475G forms puncta, enriched in the spherical shell but de-enriched in the (seemingly) hollow core. Scale bar, 10 μm. **c,** Representative images of Flag-R475G in HeLa cells stained with an anti-Flag antibody. Scale bar, 10 μm. **d,** Representative images of the indicated mClvoer3-tagged WT DDX3X or DDX3X variants and the mCherry-tagged stress granule marker G3BP1 in HeLa cells. Top, diffuse mutants. Bottom, speckle mutants. Scale bar, 10 µm. **e,** Representative images of indicated mClover3-tagged WT DDX3X or DDX3X variants and the mCherry-tagged stress granule marker G3BP1 in HeLa cells upon arsenite treatment (500 µM, 1 hr). Scale bar, 10 µm.

**Extended Data Fig. 2:**
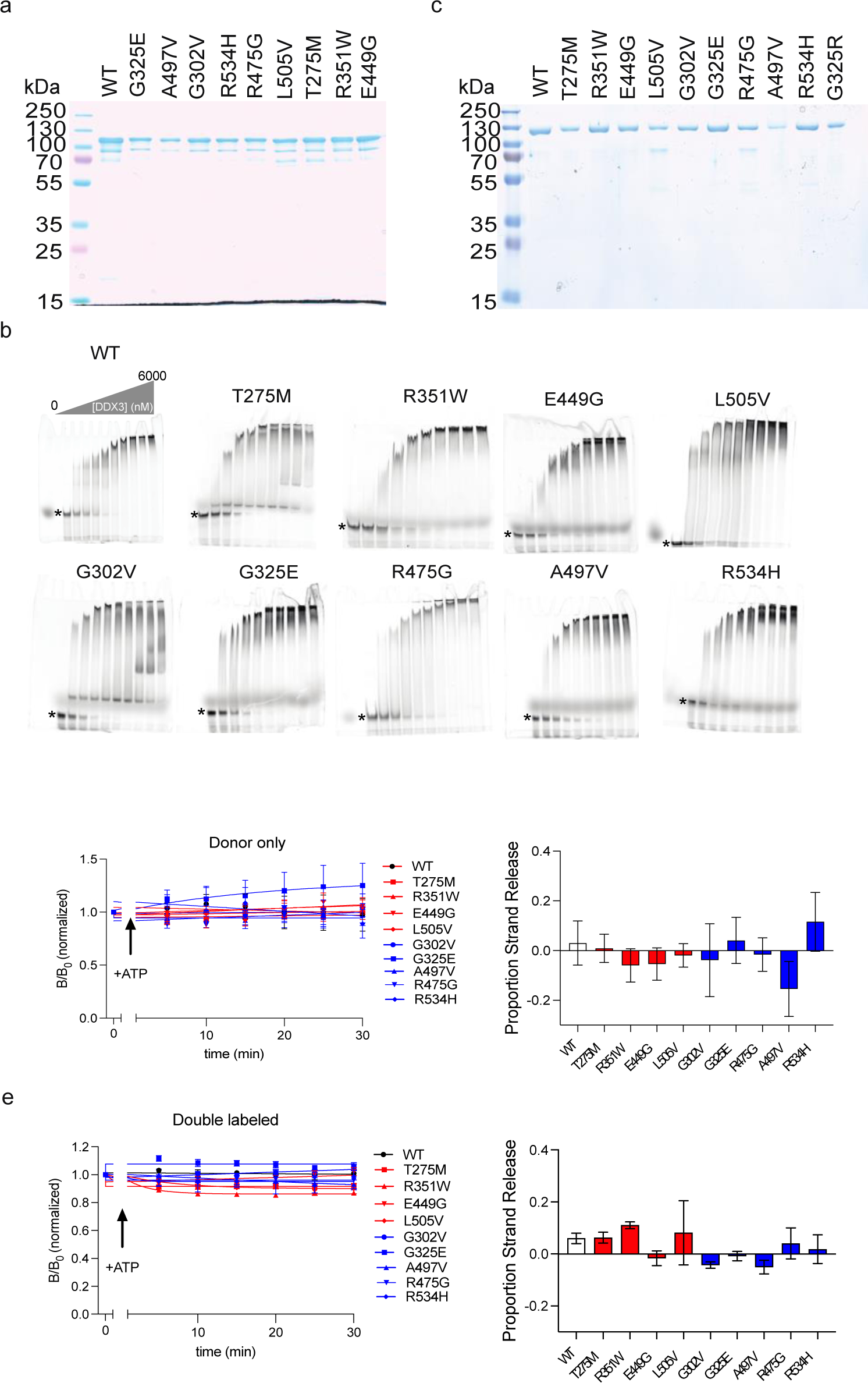
Hollow mutants are defined by their decreased ATPase and strand release activities. **a,** Coomassie-stained SDS-PAGE gel showing each purified mCherry-tagged construct. **b,** Representative EMSAs for each indicated construct. Fraction unwound was calculated for each construct by quantifying the disappearance of the unbound ssRNA (indicated by the *) at each concentration. **c,** Coomassie-stained SDS-PAGE gel showing each purified MBP-tagged construct. **d,** Left, time courses tracing the B/B_0_ (or fraction anisotropic) long strand RNA over time in the presence of 1mM ATP and MBP-tagged WT DDX3X or the indicated mutants. Right, proportion strand release graphed for each protein. **e,** Left, time courses tracing the B/B_0_ (or fraction anisotropic) duplex strand RNA over time in the presence of 1mM ATP and MBP-tagged WT DDX3X or the indicated mutants. Right, proportion strand release graphed for each protein.

**Extended Data Fig. 3:**
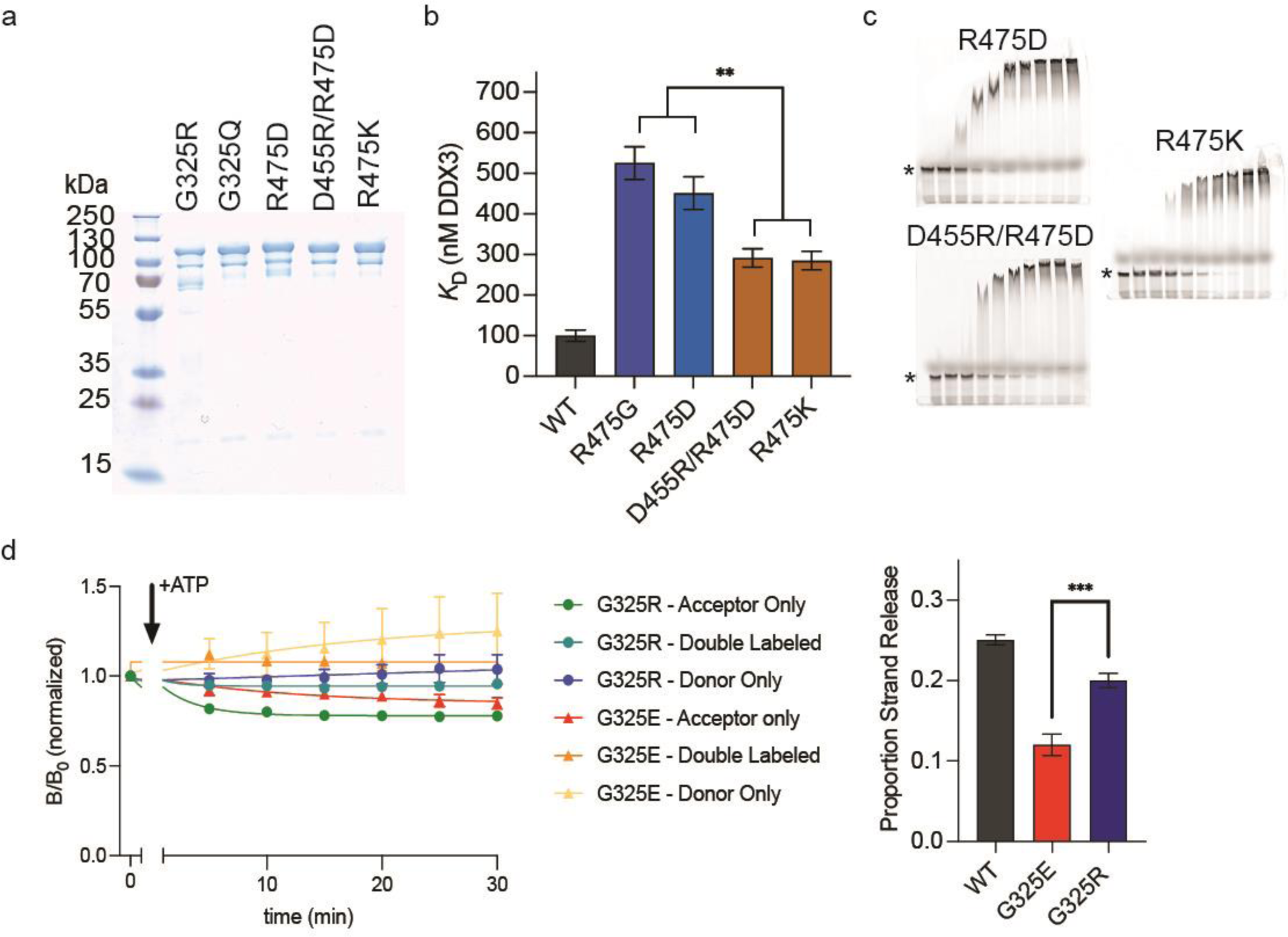
Mechanistic studies reveal how DDX3X mutants interfere with catalysis. **a,** Coomassie-stained SDS-PAGE gel showing each purified mCherry-tagged rescue mutant construct. **b,** Summarized *K_D_*values for the indicated construct, calculated via EMSA using mCherry-tagged protein in D. Values represent mean ± s.e.m., n = 3. **c,** Representative EMSAs for each indicated construct. Fraction unwound was calculated for each construct by quantifying the disappearance of the unbound ssRNA (indicated by the *) at each concentration. **d,** Changes in anisotropy signal as in Figure 2D measured for WT, G325E, and G325R.

**Extended Data Fig. 4:**
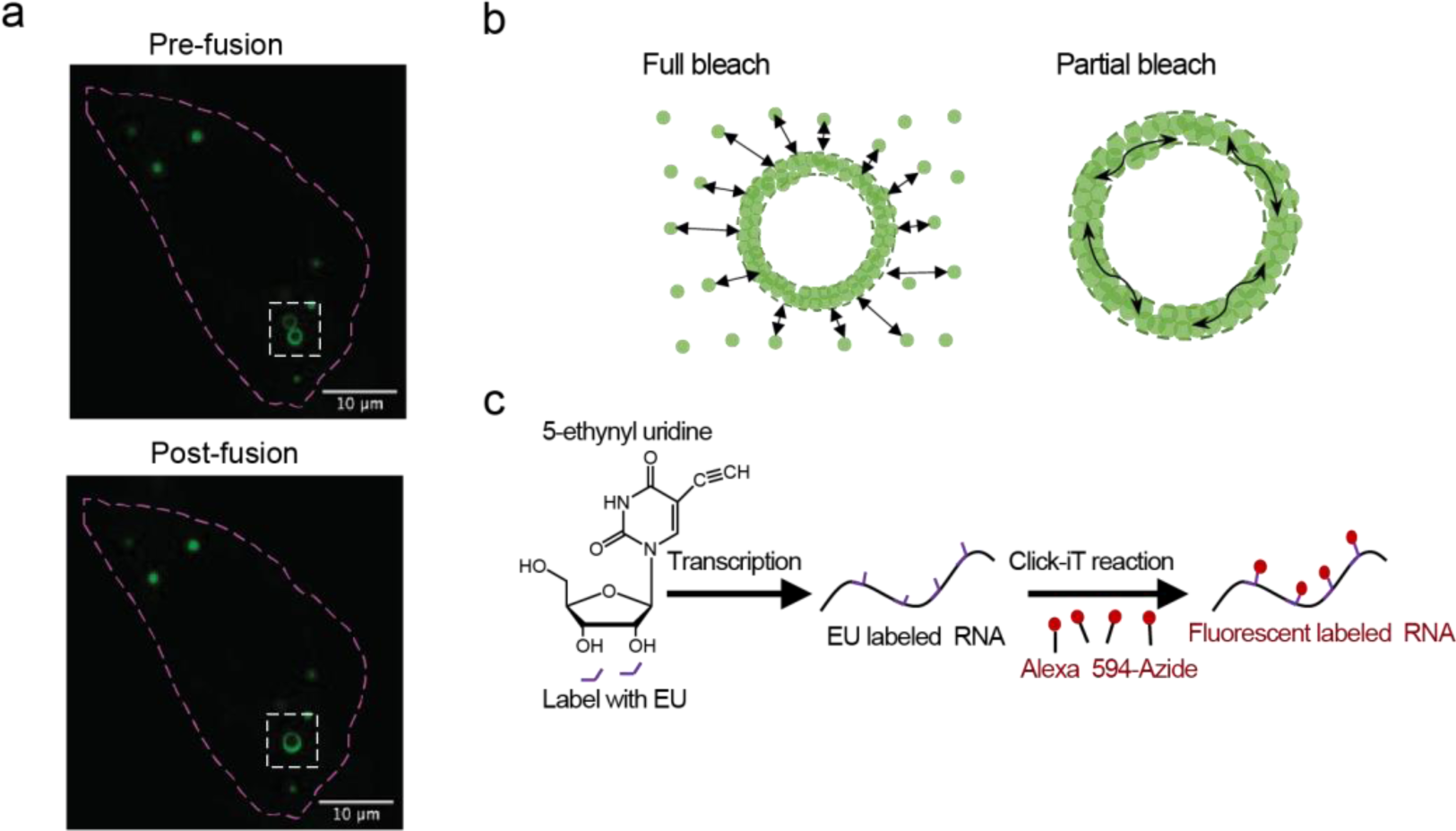
RNA-binding affinity differentiates the inter- and intramolecular dynamics of DDX3X hollow puncta. **a,** Representative still images of live-cell imaging showing the fusion of two hollow puncta of G325E-mClover3 in HeLa cells. Scale bar, 10 µm. **b,** Schematic of full bleach and partial bleach FRAP assays. Full bleach experiments were used to test for molecular exchange between the cytoplasm and hollow condensates (intermolecular). Partial bleach was used to test for the mobility of molecules within the hollow condensates (intramolecular). **c,** Schematic of the EU-labeling experiments.

**Extended Data Fig. 5:**
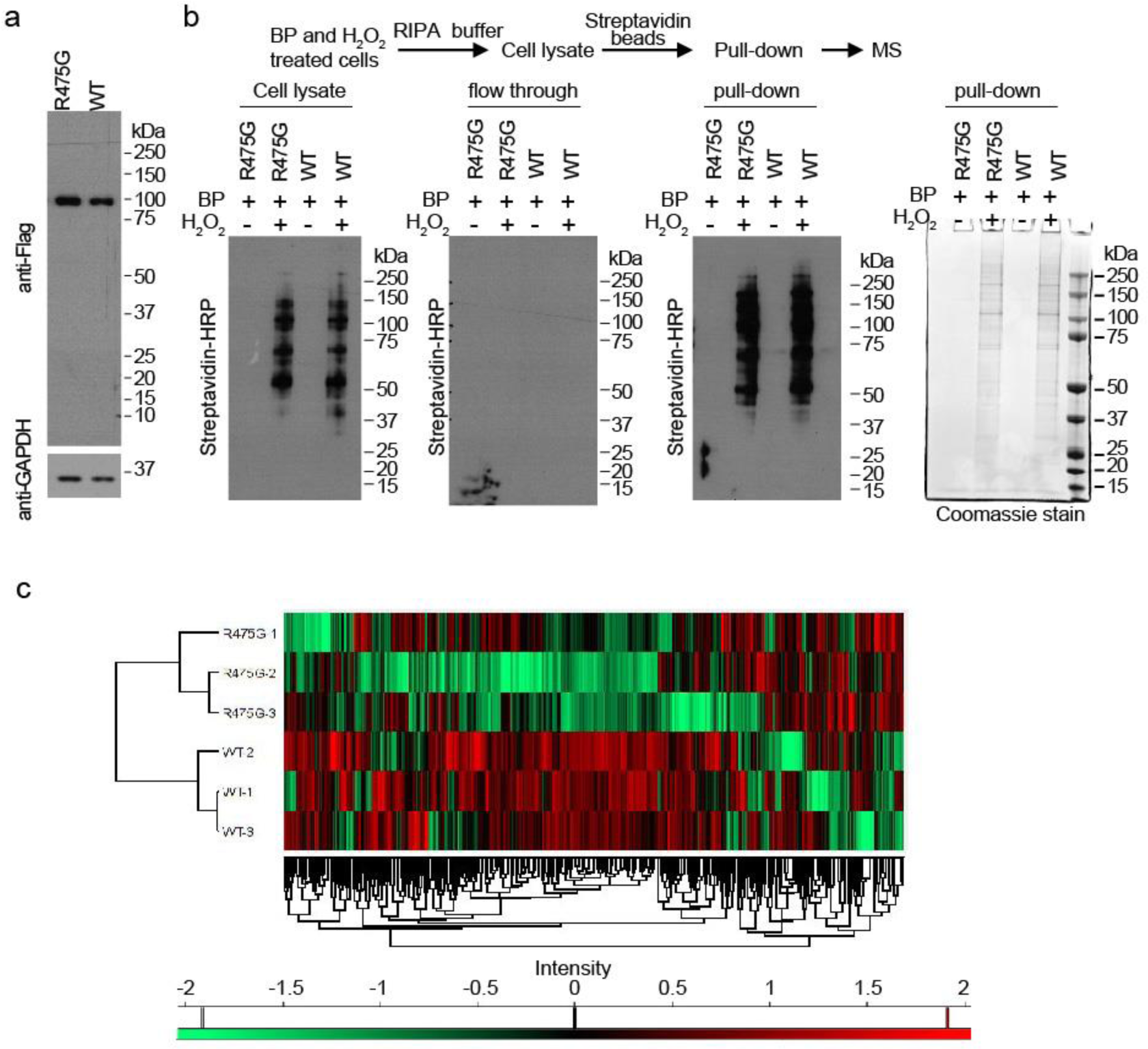
R475G hollow condensates enrich proteins in various signaling pathways and deplete translation machinery. **a,** Western blots showing the comparable expression of APEX2-tagged WT and R475G DDX3X in HEK293T cells. **b,** Top panel: workflow of the APEX2-MS sample preparation. Bottom panel: western blots and Coomassie stain showing the number of labeled proteins in the cell lysates, flow through, and pull-down fractions of streptavidin pull-down. BP: biotin phenol. **c,** Unsupervised hierarchical cluster analyses of protein abundances from quantitative APEX2-MS analyses of WT and R475G biological triplicates.

**Extended Data Fig. 6:**
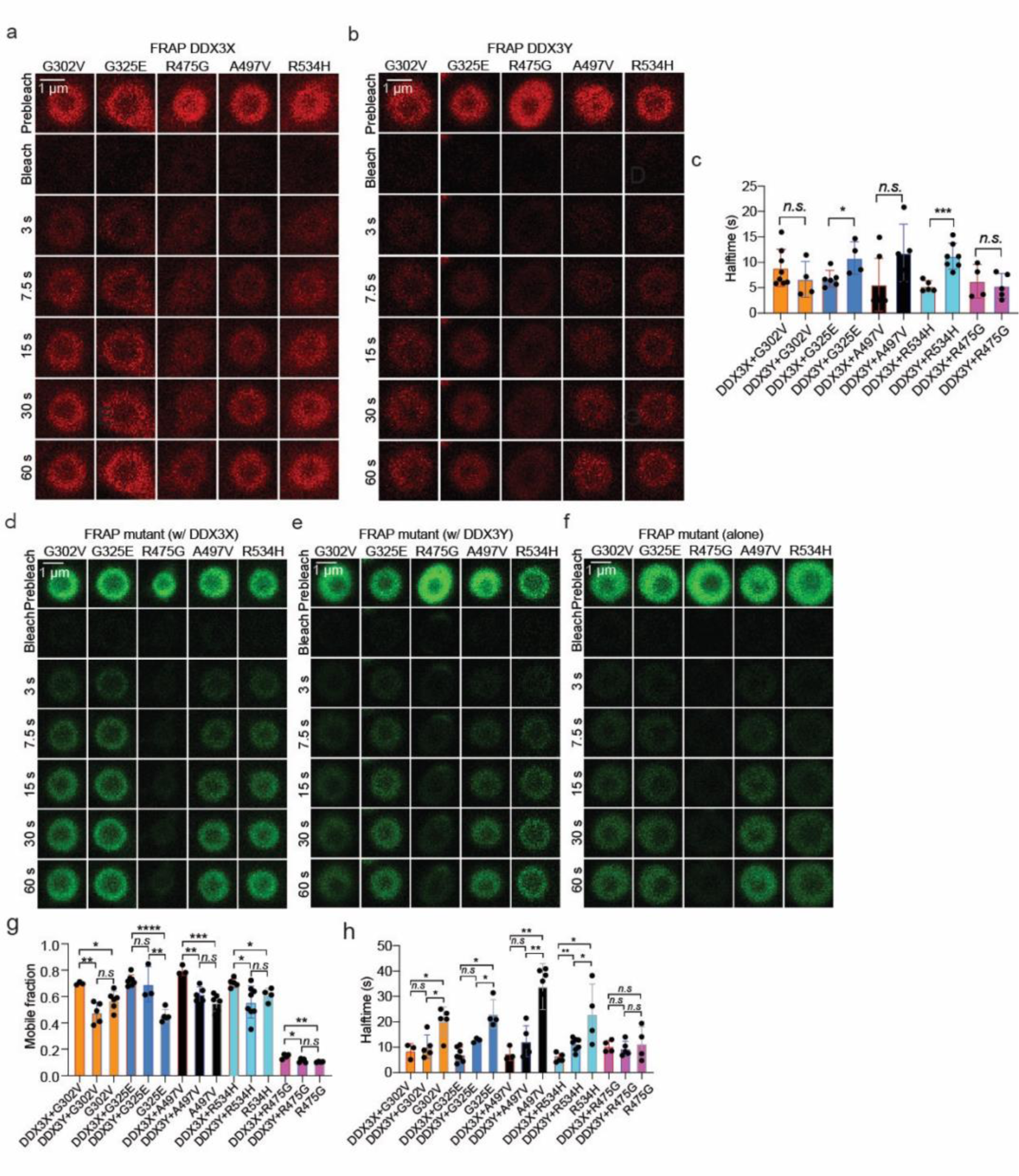
Mutant hollow condensates trap WT DDX3X and DDX3Y. **a,** Time-lapsed images of photobleached mCherry-tagged WT DDX3X trapped in the cytoplasmic hollow condensates formed by mClover3-tagged DDX3X hollow variants in HeLa cells from the cellular FRAP experiments. Scale bar, 1 µm. **b,** Time-lapsed images of photobleached mCherry-tagged WT DDX3Y trapped in the cytoplasmic hollow condensates formed by mClover3-tagged DDX3X hollow variants from the cellular FRAP experiments. Scale bar, 1 µm. **c,** Comparison of FRAP recovery halftimes of DDX3X and DDX3Y trapped in the cytoplasmic hollow condensates formed by DDX3X variants. Halftime was calculated as described in the methods. Values represent means ± s.d. from at least three biologically independent experiments. Significance was calculated using a two-tailed t-test. *p<0.05; \*\*\**p*<0.001; *n.s.* means p>0.05. **d,** Time-lapsed images of photobleached mClover3-tagged variant DDX3X in the cytoplasmic hollow co-condensates with mCherry-tagged WT DDX3X in HeLa cells from the cellular FRAP experiments. Scale bar, 1 µm. **e,** Time-lapsed images of photobleached mClover3-tagged variant DDX3X in the cytoplasmic hollow co-condensates with mCherry-tagged DDX3Y in HeLa cells from the cellular FRAP experiments. Scale bar, 1 µm. **f,** Time-lapsed images of photobleached mClover3-tagged variant DDX3X in the cytoplasmic hollow condensates when expressed alone in HeLa cells from the cellular FRAP experiments. Scale bar, 1 µm. **g,** Comparison of the FRAP mobile fractions of the indicated mClover3-tagged DDX3X variant in cytoplasmic hollow condensates either alone, with mCherry-tagged WT DDX3X or mCherry-tagged WT DDX3Y. The mobile fraction was calculated as described in the methods. Values represent means ± s.d. from at least three biologically independent experiments. Significance was calculated using a two-tailed t-test. *n.s.* means *p*>0.05; \**p*<0.05; \*\**p*<0.01; \*\*\**p*<0.001; \*\*\*\**p*<0.0001. **h,** Comparison of the FRAP recovery halftimes of the indicated mClover3-tagged DDX3X variant in cytoplasmic hollow condensates either alone, with mCherry-tagged WT DDX3X or mCherry-tagged WT DDX3Y. Halftime was calculated as described in the methods. Values represent means ± s.d. from at least three biologically independent experiments. Significance was calculated using a two-tailed t-test. *p<0.05; \*\*\**p*<0.001; *n.s.* means p>0.05.

